# Effect of reduced genomic representation on using runs of homozygosity for inbreeding characterization

**DOI:** 10.1101/2022.08.26.505374

**Authors:** Eléonore Lavanchy, Jérôme Goudet

## Abstract

Runs of homozygosity (ROHs) are proxy for genomic Identical-by-Descent segments and are increasingly used to measure individual inbreeding. ROHs analyses are mostly carried out on SNPs-arrays and whole-genome-sequencing data. Softwares recurrently used for their detection usually assume that genomic positions which have not been genotyped are non-variant. This might be true for whole-genome-sequencing data, but not for reduced genomic representations and can lead to spurious ROHs detection. We simulated the outputs of whole-genome-sequencing, two SNP-arrays and RAD-sequencing for three populations with different sizes. We compare the results of ROHs calling with two softwares: *PLINK* and *RZooRoH*. We demonstrate that to obtain meaningful estimates of inbreeding coefficients, *RZooRoH* requires fraction of genome seven times smaller compared to *PLINK*. When the SNP density is above 20 SNPs/Mb for *PLINK* and 3 SNPs/Mb for *RZooRoH*, ranks of ROHs-based inbreeding coefficients are conserved among individuals. With reduced genomic representations, ROHs distributions are consistently biased towards an underestimation of the total numbers of small and an overestimation of the total numbers of large ROHs, except for *RZooRoH* and high-density SNPs-arrays. We conclude that both ROHs-based inbreeding coefficients and ROHs distributions exact quantification are highly dependent on the fraction of genome sequenced and should thus be treated with caution. However, relative inbreeding estimates, such as comparison between individuals or populations, are reliable with reduced genomic representations providing that the fraction of genome sequenced is large enough. Consequently, we advise researchers working with reduced genomic data to use SNPs-independent measures or model-based ROHs calling methods for inbreeding estimations.

## INTRODUCTION

Inbreeding is defined as the result of mating between relatives and has been observed across many different taxa including humans (Bittles & Black, 2010; Ceballos et al., 2018; Leutenegger et al., 2011), livestock (Forutan et al., 2018; Howard et al., 2017; Kim et al., 2013; Peripolli et al., 2017, 2018), wild animal populations (Åkesson et al., 2016; Huisman et al., 2016; Kardos et al., 2018; Keller & Waller, 2002) and plants (Kariyat & Stephenson, 2019; Keller & Waller, 2002; Menges, 1991; Zhang et al., 2019). Its quantification as well as the understanding of its deleterious consequences – called inbreeding depression – are central in many areas of biology, from human genetics to conservation biology (Keller & Waller, 2002). Indeed, increase in genome autozygosity has been associated with humans diseases, such as schizophrenia (Keller et al., 2012; Lencz et al., 2007) and Alzheimer’s disease (Ghani et al., 2015; Nalls et al., 2009) as well as fitness costs in both animal (Åkesson et al., 2016; Bjelland et al., 2013; Huisman et al., 2016; Parland et al., 2007) and plants (Menges, 1991; Zhang et al., 2019).

Individual levels of inbreeding are quantified by estimating individuals inbreeding coefficients (*F*). Traditionally, inbreeding was measured counting the size and number of loops in pedigrees (*F*_*PED*_) (Wright, 1922) a method that has several downsides: i) it estimates the expected inbreeding coefficient which can differ from the realized coefficient due to recombination stochasticity and mendelian segregation (Carothers et al., 2006; Franklin, 1977; Hill & Weir, 2011); ii) it assumes all founders of the pedigree are unrelated and non-inbred; iii) pedigrees must be correctly recorded which is extremely difficult in wild populations, although genetic data might be used to (re)construct correct links (Huisman, 2017; Jones & Wang, 2010). With the advancements in high throughput sequencing technologies it became possible to estimate with sufficient accuracy genomic-based inbreeding coefficients, and several studies have shown molecular estimates to be more accurate than pedigree-based estimates (Alemu et al., 2020; Curik et al., 2014; Kardos et al., 2015; Keller et al., 2011; Wang, 2016). Many different genomic-based inbreeding coefficients have then been proposed, such as *F*_*HOM*_ (Chang et al., 2015; Purcell et al., 2007), *F*_*AS*_ (Weir & Goudet, 2017), *F*_*UNI*_ and *F*_*GRM*_ (both described in Yang et al., 2011) but there is still no consensus on which is the most accurate (Alemu et al., 2020; Caballero et al., 2020; Goudet et al., 2018; Nietlisbach et al., 2019; Yengo et al., 2017; Zhang et al., 2022). These estimates quantify average excess SNP homozygosity or correlation between uniting gametes and treat all SNPs independently. However, parents transmit DNA to their offspring in large chromosomal segments rather than each base independently. Consequently, it has been suggested that measures of inbreeding coefficients should be based on Identical-by-Descent (IBD) genomic segments rather than individual SNPs (McQuillan et al., 2008). Hence, a new inbreeding coefficient *F*_*ROH*_, was proposed by McQuillan et al., (2008).

This novel coefficient is based on Runs of Homozygosity (ROHs), long consecutive homozygous segments (Ceballos et al., 2018). ROHs arise when two IBD segments are brought together in an individual as a result of parents’ co-ancestry (Broman & Weber, 1999; Ceballos et al., 2018). ROHs were first described by (Broman & Weber, 1999) and shown to be ubiquitous in humans (Ceballos et al., 2018; Gibson et al., 2006; Pemberton et al., 2012) and across many different taxa (Kardos et al., 2018; Liu et al., 2020; Saremi et al., 2019). The ROHs-based inbreeding coefficient *F*_*ROH*_ is traditionally calculated as the proportion of the genome within ROHs and several studies demonstrated that it was a reliable estimator of inbreeding (Alemu et al., 2020; Caballero et al., 2020; Nietlisbach et al., 2019). In addition to quantifying inbreeding, ROHs distributions (i.e. lengths and numbers) can also inform about a population’s past demography and history (Bosse et al., 2012; Ceballos et al., 2018; Kirin et al., 2010; Nothnagel et al., 2010; Pemberton et al., 2012): long ROHs reflect recent inbreeding events while smaller ROHs indicate more distant inbreeding and, if in high proportion, a history of small effective population size. Besides informing about inbreeding, ROHs can be used for identifying rare deleterious recessive variants responsible for deleterious phenotypes by homozygosity mapping which in short compares sick and healthy people ROHs islands (Alkuraya, 2013; Hildebrandt et al., 2009; Keller et al., 2012; Lencz et al., 2007; Wang et al., 2009).

Two different methods for ROHs detection are recurrently used in the literature: observation and model-based approaches (Ceballos et al., 2018). The most common method is a fast observation-based method (Ceballos et al., 2018) implemented in *PLINK* (Chang et al., 2015; Purcell et al., 2007). It makes use of a sliding window to identify continuous homozygous stretches with a minimum size defined by the user. Model-based approaches, such as those implemented in *RZooRoH* (Bertrand et al., 2019; Druet & Gautier, 2017), *BEAGLE* (Browning & Browning, 2010) and *BCFTools* (Narasimhan et al., 2016) rely on Hidden Markov Models (HMM) and do not require a minimum threshold on ROHs length. HMM methods are computationally demanding (Ceballos et al., 2018) and a previous study suggested than *PLINK* outperformed HMM methods both in terms computation time and ROHs detection accuracy with simulated whole-genome-sequencing (WGS) data (Howrigan et al., 2011).

Observation-based approaches were designed for WGS data and assume that the region between two SNPs are entirely homozygous. However, many studies performing ROHs analyses with *PLINK* use reduced genomic representation techniques: ROHs have been very often called with SNP arrays (Bjelland et al., 2013; Bosse et al., 2012; de Jong et al., 2020; Forutan et al., 2018) where specific SNPs, chosen based on their position, effect on phenotype or Minor Allele Frequency (MAF), are targeted and sequenced. ROHs have also been called with Restriction-Site Associated DNA Sequencing (RAD-sequencing) (Grossen et al., 2018; Mueller et al., 2022) data, by cutting the genome near enzymes cutting sites and then selecting and sequencing fragments based on their size. With both SNP arrays and RAD sequencing, only a (small) fraction of the genome is sequenced resulting in a partial representation of the total polymorphism. Since *PLINK* assumes that genomic positions not included in the SNPs set are nonvariant, we expect that they will falsely consider non-sequenced heterozygous loci as homozygous which can lead to spurious ROHs detection. On the contrary, the HMM approach from Leutenegger et al. (2003), which models the genome as a mosaic of IBD and non-IBD segments and from which most current model-based approaches follow, was initially developed for SNPs arrays. These models do not treat non-sequenced genomic regions as homozygous but rather as missing data. However, model-based approaches are rarely used for ROHs analyses with reduced genomic data. In addition, no precise benchmarking with large sample size has been performed on comparing how the different ROHs calling methods behave with these reduced genomic data compared to WGS data and precise guidelines such as which method is suitable with which data are missing.

In addition to the fraction of genome captured, we hypothesize that the effective size and level of polymorphism in a population might also affect the capacity of the different methods to accurately detect ROHs. On one hand, small and inbred populations will tend to harbour higher numbers of long ROHs easier to accurately detect with reduced representations as the missing positions are more likely to be homozygous. In addition, larger populations will tend to harbour many small ROHs (Ceballos et al., 2018; Kirin et al., 2010) harder to correctly identify when only a fraction of the polymorphism is available since these small ROHs require high number of nearby SNPs to reach the minimum density threshold for ROH detection (Kardos et al., 2015; Sole et al., 2017; Zhang et al., 2015). On the other hand, larger populations will harbor higher levels of polymorphism and thus higher numbers of SNPs resulting in an increased SNP density for the same fraction of genome sequenced.

Here, we use simulated data to compare the performance of ROHs calling from WGS – considered our gold standard – and two reduced genomic representations – SNP arrays and RAD-sequencing – as well as the two ROHs calling methods implemented in *PLINK* and *RZooRoH* in both a small and a large populations. We hypothesize that the quality of ROHs detection will depend on the fraction of genome covered with the reduced sequencing method. In addition, since model-based approaches take into account the distance between each SNP, we predict that they will perform better when dealing with sparse data (Druet & Gautier, 2017). We show that both ROHs detection methods can be used to correctly estimate ROHs-based inbreeding coefficient *F*_*ROH*_ with SNP arrays and RAD-sequencing providing that a sufficient proportion of the genome has been sequenced. This proportion varied between ROHs calling methods and population sizes: model-based method implemented in *RZooRoH* as well as the large population require a substantially smaller fraction of the genome to obtain correct *F*_*ROH*_ estimates. However, while inbreeding coefficients based on ROHs can be correctly estimated, ROHs distributions are almost always biased with reduced genomic representations as the number of small ROHs is constantly underestimated while the number of large ROHs is constantly overestimated.

## MATERIAL & METHODS

### Simulations

A general workflow of the study can be found in figure 1 and additional details about the simulations and analyses performed can be found in supplementary material. To investigate the effect of population size on ROHs calling with reduced genomic representation, we simulated two hermaphroditic populations (N = 1,000 and N = 10,000) using SLiM3, a forward-in-time individual-based simulation software (Haller & Messer, 2019). We used a ‘non-Wright Fisher’ model which included non-fixed population sizes and overlapping generations to increase the chances of related mating (Haller & Messer, 2019). The population size was regulated via a patch carrying capacity where individuals were removed based on their overall fitness at the end of each simulation cycle. Individuals’ fitness decreased based on their age which varied between 0 and 3, with older individuals having higher probabilities to die. Individuals were able to reproduce from the age of 1 and selfing was not allowed. For each individual, its mate was chosen among the other individuals based first on their age (with older individuals less likely to be chosen) and second on their pedigree-based coancestry with the reproducing individual (related individuals had higher chances to be chosen than non-related ones). This resulted in a population mostly practicing random mating but ensured that some inbreeding would occur at each generation. We simulated ten replicates for each of the two population sizes and each simulation lasted for 1,000 reproductive cycles. We used a human-like genetic map with a non-homogenous recombination rate along the genome simulated with *FREGENE* as described in Chadeau-Hyam et al. (2008). Individuals from both populations carried 30 chromosomes each 100 Mb long. The burn-in were performed via *recapitation* in *msprime* (Kelleher et al., 2016) as suggested by Haller et al. (2019). All mutations were added at the end of the simulation (after the burn-in) based on a human-like mutation rate of 2.5^*e*-8^ (Nachman & Crowell, 2000) as suggested by Haller *et al*. (2019) and Kelleher *et al*. (2018).

**Figure 1:**
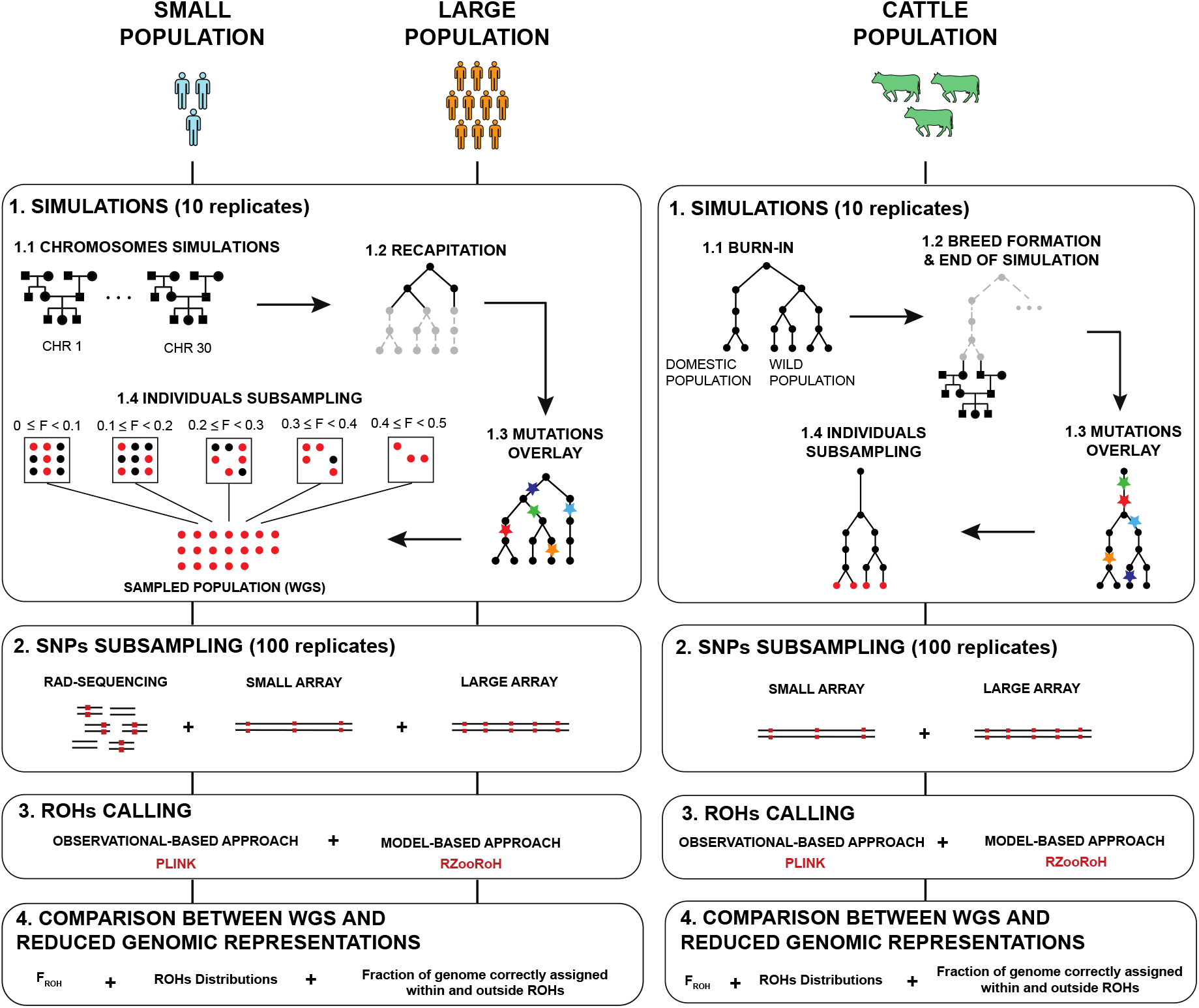
General workflow of the study. Simulations were first performed in SLiM3^1^ where each chromosome of the same population shared the same pedigree; burn-in, recapitation and mutation overlay were then performed in msprime^2,3^. SNPs subsampling was performed with bedtools^4^ for RAD-sequencing and home-made python script for both arrays. ROHs were called both with PLINK and RZooRoH and F_ROH_ was estimated as the fraction of the genome within ROHs^5^. ROHs distributions were divided into six length classes and fraction of genomes correctly and incorrectly assigned within and outside ROHs were estimated as the overlap between ROHs obtained with WGS data and ROHs obtained with reduced genomic representation data.

At the end of the simulation, we performed a random stratified sampling to ensure that the individuals used in subsequent analyses would cover the entire range of inbreeding. We are aware that this scheme is rare and hard to apply empirically but it allowed us to investigate whether the entire spectra of inbreeding was correctly detected. Whenever possible, we subsampled twenty individuals with *F*_*PED*_ between 0 and 0.1, twenty individuals with *F*_*PED*_ between 0.1 and 0.2, twenty individuals with *F*_*PED*_ between 0.2 and 0.3, twenty individuals with *F*_*PED*_ between 0.3 and 0.4 and finally twenty individuals with *F*_*PED*_ between 0.4 and 0.5. Average (±SD) number of sampled individuals per replicate were 87.3 ± 5.03 for the small population and 67.40 ± 4.09 for the large population. The lower number of individuals subsampled in the large populations are because larger populations contained fewer individuals with high inbreeding coefficients. The mean (±SD) number of SNPs per simulation was 1.6e^06^ ± 2.0e^04^ SNPs for the small population and 1.6e^07^ ± 1.7e^05^ SNPs for the large population.

### SNPs subsampling

In order to investigate the effect of reduced genomic representations on ROHs detection, we mimicked different sequencing techniques by subsampling SNPs from whole-genome data. We simulated both RAD-Sequencing and two SNP arrays of different sizes.

RAD-sequencing uses restriction enzymes to digest the genome in small fragments, which are then selected on size and sequenced (Andrews et al., 2016). Consequently, these fragments are not homogenously distributed along the genome. For this purpose, we randomly selected 500 base-pair (bp) fragments (Andrews et al., 2016) using *bedtools v2*.*29* (Quinlan, 2014). Afterwards, SNPs within these windows were subsampled using *--bed* function from *VCFTools* (Danecek et al., 2011). Given that the proportion of the genome sequenced with RAD-sequencing vary greatly depending on the organism and on the restriction enzymes used, we varied this number of fragments so that they covered between 0.05% and 15% of the genome, and between 0.002% and 1%, for the small and the large populations respectively. We performed 100 replicates for each subsampling percentage.

To simulate SNP array sequencing, we mimicked two arrays initially developed for cattle and widely used for ROHs analyses: the Illumina BovineSNP 50 beadchip (∼ 50’000 SNPs) hereafter “small array” - and the Illumina BovineHD BeadChip – (∼ 777’000 SNPs) hereafter “large array”. Common features of both arrays are the homogenous distances between SNPs and the focus on common SNPs, hence, we first filtered our WGS data on MAF 5%. We then selected windows with size corresponding to the median distances between SNPs in the real arrays – 40kb for the small array and 3kb for the large array – and selected the SNP with higher MAF within each window (if at least one SNP was present). Please note that we use the term ‘small array’ for what is usually considered as a medium-density array in the cattle literature.

### ROHs calling

We compared two methods for ROHs detection to investigate whether we observe a difference in their capacity to handle reduced genomic data. We chose one observational-based approach – the *--homozyg* method, implemented in *PLINK* (Chang et al., 2015; Purcell et al., 2007) – and one model-based approach – the *RZooRoH* method, implemented as a R package (Bertrand et al., 2019; Druet & Gautier, 2017).

For ROHs detection with *PLINK*, we used the default values for all parameters except one: we authorized zero heterozygous SNPs per windows (*--homozyg-window-het = 0*) as we have simulated data and there is no uncertainty about the call of each variant. We chose a minimum size of 100KB for a ROH to be detected. These parameters were consistent for every replicate in each subsampling method.

We then called ROHs with the *RZooRoH* package with a three homozygosity-by-descent (HBD) classes model with rates equals 8, 16 and 32 for the HBD classes and 256 for the non-HBD class. These classes correspond to different inbreeding events ages and, for each class, the rate corresponds to the expected number of generations since the inbreeding event divided by 2. We chose a model with few classes due to computational constrains and because it resulted in inbreeding coefficients values similar to the ones obtained with F_*PED*_ and with PLINK minimum ROHs size 100KB with WGS data (FIG S1).

In order to verify the performance of *RZooRoH* and *PLINK*, we compared the results of ROHs calling with WGS data and both softwares. Overall *RZooRoH* and *PLINK* led to similar results and ranks of inbreeding among individuals were mostly conserved (FIG S1). However, we observe slight differences: first, we show that total length of all ROHs were higher with *RZooRoH* compared to *PLINK* (FIG S1). We also show that *F*_*ROH PLINK*_ is closer to the true proportion of IBD segments up to 100 generations ago while *F*_*ROH RZooRoH*_ is closer to the true proportion of IBD segments older than 100 generations ago (more details in supplementary material, FIG S2, S3 & S4).

### Statistical analyses

For *PLINK*, we estimated individual ROHs-based inbreeding coefficient (*F*_*ROH*_) as the proportion of the genome within ROHs: 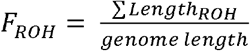 (McQuillan et al., 2008). For *RZooRoH*, we estimated *F*_*ROH*_ as the average HBD probabilities (the @realized) as suggested in the *RZooRoH* documentation. We then compared *F*_*ROH*_ estimated with WGS that we consider as our ‘gold standard’ and the reduced-representation sequencing technologies with Pearson correlations and linear regression.

Since ROHs distribution can inform about the population history (Ceballos et al., 2018), we divided ROHs into six length classes following Kirin et al., (2010): i) between 100Kb and 2Mb [i.e. between 0.1cM and 2cM], ii) between 2Mb and 4Mb [i.e. between 2cM and 4cM], iii) between 4Mb and 6Mb [i.e. between 4cM and 6cM], iv) between 6Mb and 8Mb [i.e. between 6cM and 8cM], v) between 8Mb and 16Mb [i.e. between 8cM and 16cM] and, vi) larger than 16Mb [i.e. larger than 16cM]. ROHs distributions are represented as the mean total length per individual among simulation and subsampling replicates. This length classes are traditionally used with *PLINK* since *RZooRoH* has its own ROHs distributions estimation based on the rates chosen when constructing the model. However, we chose to estimate these length classes for both detection methods for the sake of comparison and because *RZooRoH* relies on the average probabilities of belonging to any length class for each SNP which does not result in distributions *per se*. We then compared the distributions between WGS (our ‘gold standard’) and the other sequencing techniques.

We use four metrics to evaluate the accuracy of ROHs detection for each subsampling technique: i) the fraction of genome correctly assigned within ROHs (true-ROHs) i.e. ROHs which were detected with both WGS and the reduced genomic representation, ii) the fraction of genome correctly assigned outside ROHs (true-non-ROHs) i.e. genomic regions which were not classified as ROHs with neither WGS nor the reduced genomic representation, iii) the fraction of genome inappropriately assigned within ROHs (false-ROHs) i.e. genomic regions which were not classified as ROHs with WGS but detected as ROHs with the reduced genomic representation and finally iv) the fraction of genome inappropriately assigned outside ROHs (false-non-ROHs) i.e. ROHs which were detected with WGS but not assigned as ROHs with the reduced genomic representation. We compared ROHs calling between WGS and reduced representation for every individual in each replicate, subsampling method and simulation. We then averaged individual’s fractions among simulation and subsampling replicates to obtain one measure per subsampling event.

### Additional Simulations

We also performed an additional batch of simulations based on a real, 57 years deep, cattle pedigree from Walloon beef cattle. We used a genetic map estimated from male Holstein cattle by Qanbari & Wittenburg (2020). In the simulation, a domestic population (Ne = 1500) got separated from a large wild population (Ne = 50,000) 10,000 generations ago with a migration rate of 3^*e*-5^ (Frantz et al., 2020). To mimic the strong selective pressure which occurred during breed formation 200 generations ago and which resulted in high levels of inbreeding (Frantz et al., 2020), 200 individuals were randomly selected from the domestic population and used as founders the rest of the simulation. The remaining 200 generations were then simulated in *SLiM3* from these 200 founders. As the real pedigree was only covering the last 57 years, a first round of simulations was run to obtain 200 generations-deep simulated pedigree, which was then used to complete the real pedigree by assigning a genealogy from the simulated pedigree to each founder from the real pedigree. At the end, only the individuals from the real pedigree were kept for the analyses.

Since we showed with the first batch of simulations that the accuracy of ROHs detection with RAD-sequencing depends on the proportion of genome subsampled, we only mimicked SNP-Arrays-like subsampling for these simulations. We did so as described previously. However, at the end of the sliding windows process, we obtained a lower number of SNPs than expected as some windows did not contain any SNPs. Consequently, additional SNPs were chosen randomly to account for empty windows and to reach the same number of SNPs as in real arrays.

## RESULTS

### F^ROH^

We used simulated genomes to investigate the influence of different sequencing techniques, ROHs calling methods and, population size on ROHs detection. Figure 2 shows *F*_*ROH*_ estimated using reduced representations sequencing techniques data (*F*_*ROH RAD*_ and *F*_*ROH ARRAY*_) compared to *F*_*ROH*_ estimated using WGS data (*F*_*ROH WGS*_ our ‘gold standard’). Points closer to the black equality line (x = y) indicate that *F*_*ROH RAD*_ and *F*_*ROH ARRAY*_ are similar to individual *F*_*ROH WGS*_; points below the equality line indicate underestimated values and points above the equality line overestimated values. Figure 2 shows *F*_*ROH*_ can be correctly estimated with reduced genomic representations providing that a sufficient fraction of the genetic variation is captured. With RAD-sequencing (FIG 2, panels A and B), *PLINK* (FIG 2, panel A) required roughly seven times more data than *RZooRoH* (FIG 2, panel B) for *F*_*ROH RAD*_ to become consistent with *F*_*ROH WGS*_ (*PLINK* correctly estimated *F*_*ROH*_ and the rank in inbreeding among individuals with 10% of genome in the small and 1% in the large population, FIG 2, panel A; FIG S6, S7. *RZooRoH* correctly estimated both *F*_*ROH RAD*_ estimates and the rank of inbreeding among individuals with 1% of genome sequenced in the small and 0.125% in the large population, FIG 2, panel B; FIG S8, S9.) In addition, we show in supplementary material that *F*_*HOM*_, an estimator of inbreeding coefficient relying on the difference between the observed and expected heterozygosity under Hardy-Weinberg yielded similar results to *RZooRoH* for the same SNP densities (FIG S10, Table S1).

**Figure 2:**
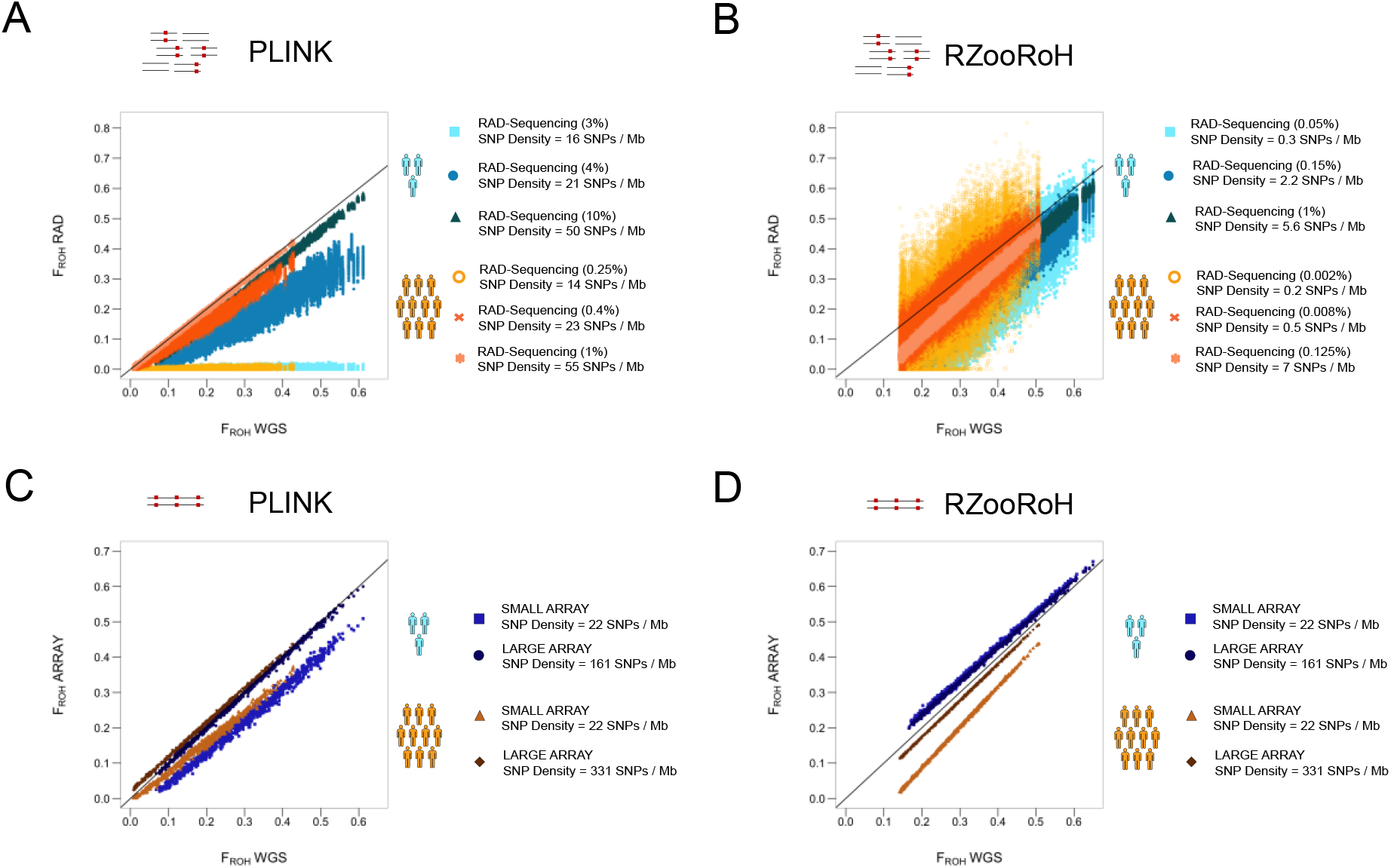
Comparison between F_ROH_ estimated with reduced genomic data on the y axis (F_ROH RAD_ & F_ROH ARRAY_) according to F_ROH_ estimated with WGS data on the x axis (F_ROH WGS_). Each point represents one individual (for one subsampling replicate within one simulation replicate). The black line represents the equality line (x = y). Blue points represent individuals from the small population and orangish from the large population. Within these two colors categories, a change in shade represents an increase in fraction of genome subsampled (indicated between the parentheses for RAD-sequencing). **A:** Subsampling was performed mimicking RAD-sequencing and ROHs were called with PLINK. **B:** Subsampling was performed mimicking RAD-sequencing and ROHs were called with RZooRoH. **C:** Subsampling was performed mimicking SNPs arrays and ROHs were called with PLINK. **D:** Subsampling was performed mimicking SNPs arrays and ROHs were called with RZooRoH.

*PLINK* did not detect ROHs with proportion of sequenced genome below 3% in the small and 0.25% in the large populations. This resulted in *F*_*ROH RAD*_ of zero for all individuals (FIG. 2, panel A). When increasing the fraction of genome represented (4% in the small and 0.4% in the large populations), *F*_*ROH RAD*_ estimates differed from zero. However, individuals *F*_*ROH RAD*_ were always underestimated but the rank of inbreeding was conserved among individuals (FIG. 2, panel A; FIG. S6, S7, Table S1). With the subsampling we used, *RZooRoH* always detected ROHs, independently of the fraction of genome sequenced we tested. However, the variance among subsampling replicates was extremely large and the rank of individuals’ inbreeding was poorly conserved for low fractions of the genome: 0.05% in the small and 0.002% and to a lesser extent 0.008% in the large populations (FIG 2, panel B; FIG S8, S9, Table S1).

Concerning SNP arrays (FIG 2, panels C and D), we see little effect of the type of array or the software used: *F*_*ROH ARRAY*_ estimates were consistent with *F*_*ROH WGS*_ estimates, except in the large population where estimates were slightly underestimated with *RZooRoH* and the small array. The rank of inbreeding was always conserved among individuals (FIG 2, panels C and D; FIG S11, S12). We observed an effect of population size: a larger population required a smaller fraction of the genome to obtain *F*_*ROH*_ estimates similar to WGS with reduced representations (FIG. 2, panels A and B; FIG. S5). Interestingly, these minimum fractions of the genome necessary to get high correlation between WGS and RAD-sequencing *F*_*ROH*_ estimation resulted in similar SNP densities in both populations (20 SNPs per Mb with *PLINK* and 3 SNPs per Mb with *RZooRoH*) suggesting that SNP density might be a key metric for assessing the accuracy of F_ROH_ estimation and the conservation among individuals inbreeding ranks (FIG S5).

### ROHs distributions

Figure 3 shows ROHs distribution among the different length classes as the mean per individual (among simulation and subsampling replicates) total ROHs length falling within each ROH length class. Horizontal black lines represent our gold standard: the mean (among simulation replicate) individual total ROHs lengths estimated with WGS for each ROH class. Barplots represent the mean (among simulation and subsampling replicate) difference between the reduced representation and WGS. Thus, barplots above the horizontal black segment indicate an overestimation while barplots below the segment an underestimation. In addition, the y axis starts at 0 indicating than no ROHs of the particular length class has been detected if the bar reaches the bottom of this axis. Compared to WGS, reduced representations generally underestimate the number of short ROHs and overestimate the number of long ROHs. As indicated by smaller barplots’ absolute heights from *RZooRoH* compared *PLINK*, this bias is more pronounced with *PLINK* than with *RZooRoH* when both methods are applied to reduced data with same densities (i.e. with both arrays) (FIG 3 panels A and C). With *PLINK*, ROHs distributions were always biased. With *RZooRoH*, RAD-sequencing always yielded biased distributions. In the large population both arrays allowed correct estimation of total lengths of ROHs larger than 2MB. The bias was stronger for the small array compared to the large array. In the small population, ROHs distributions were correctly estimated with both arrays. Interestingly, we observed a pattern for all reduced genomic representations with *PLINK:* no ROHs smaller than 4MB were detected (i.e. total length of ROHs shorter than 4MB equal 0) with the lowest fraction of genome sequenced and the small array in both populations (FIG 3 panels A and C).

**Figure 3:**
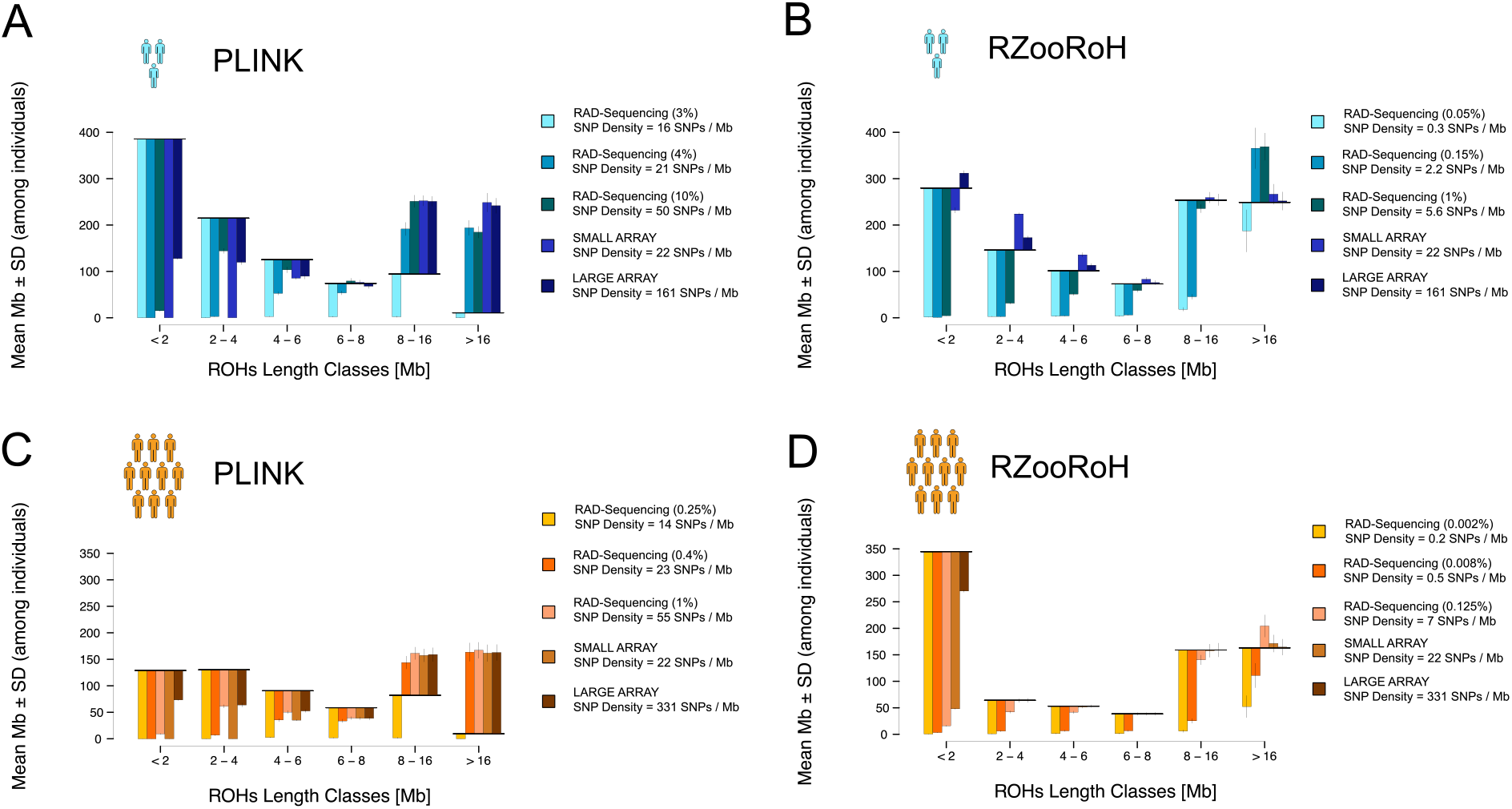
Comparison of ROHs distributions between the subsampled dataset and WGS. Black horizontal lines correspond to WGS total ROHs lengths per individuals (y axis) falling into the different ROHs length classes (x axis). Barplots show the mean (± sd) difference between WGS and each subsampling method. Barplots below the horizontal black line indicate an underestimation while barplots above the horizontal black line indicate an overestimation of the total length of ROHs estimated with the reduced genomic representation compared to WGS. Mean (and sd) are among individuals, simulation and subsampling replicates. **A:** ROHs distributions from the small population; ROHs were called with PLINK. **B:** ROHs distributions from the small population; ROHs were called with RZooRoH. **C:** ROHs distributions from the large population; ROHs were called with PLINK. **D:** ROHs distributions from the large population; ROHs were called with RZooRoH.

In order to quantify what is the minimum fraction of the polymorphism required to correctly estimate ROHs distributions, we randomly subsampled between 3 and 90 percent of the SNPs (≠ fraction of genome) and reproduced the same analyses. Figures S17-S20 show that very large proportions (above 60%) were required to reach ROHs distributions similar to WGS with *PLINK* and in both populations. Finally, we found that reduced representations tend to merges small adjacent ROHs into larger ones as illustrated in figure S21A.

### Fraction of genome assigned within and outside ROHs

Figure 4 shows the mean fraction of genome which has been correctly (true-non-ROHs and true-ROHs) and incorrectly (false-non-ROHs and false-ROHs) assigned compared to WGS (please note that the “true” fraction of genome within ROHs (i.e. with WGS) differ between both ROHs calling softwares). Concerning RAD-sequencing, the fraction of the genome correctly assigned with reduced genomic techniques was higher for *PLINK* (FIG 4 left column, panels A and C) compared to *RZooRoH* (FIG 4 right column, panels B and D) for both populations but fractions of genome sequenced with RAD-sequencing were also larger for *PLINK*. Concerning SNPs arrays, fraction of genome correctly assigned were similar between both methods in the small population but higher for *PLINK* compared to *RZooRoH* in the large population. Interestingly, the reason underlying the wrong assignment differed between both calling methods in the small population: *RZooRoH* resulted in higher fraction of genome incorrectly assigned within ROHs (false-ROHs, FIG 4 panel B) while to PLINK mostly resulted in false-non-ROHs (FIG 4 panel A). In the large population, both methods resulted mostly in false-non-ROHs (FIG 4 bottom row, panels C and D). Finally, the proportion of genome correctly assigned reached high values: minimum 90% with the small and 95% with the large array, with both populations and sequencing techniques.

**Figure 4:**
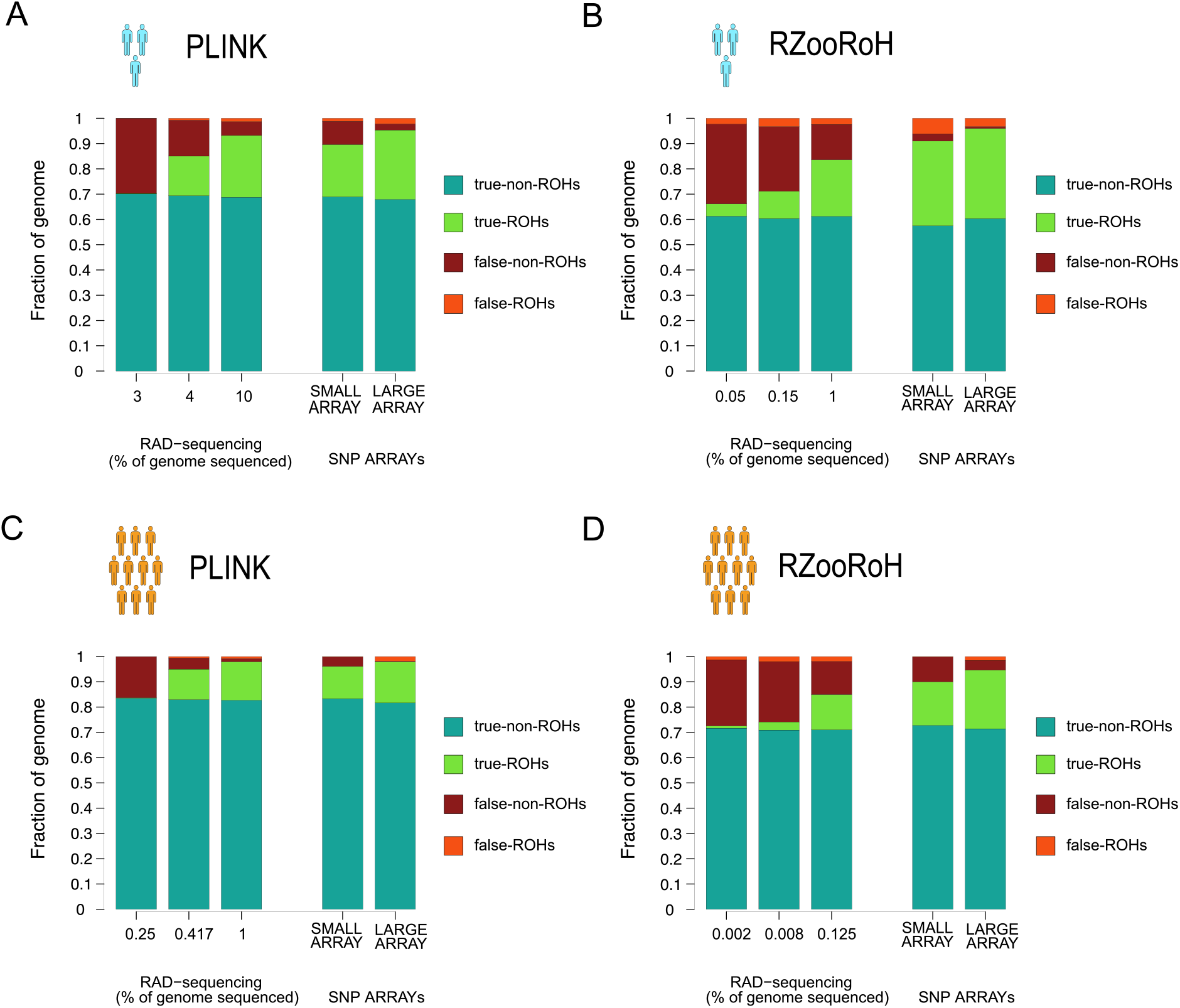
For each reduced genomic representation fraction of genome correctly assigned outside ROHs (true-non-ROHs), correctly assigned within ROHs (true-ROHs), incorrectly assigned outside ROHs (false-non-ROHs) and incorrectly assigned within ROHs (false-ROHs) are represented in regards to WGS data. Note that WGS ROHs results (and thus maximum fractions of genome correctly assigned within and outside ROHs) differ between PLINK and RZooRoH. Values are averaged among individuals and both simulation and subsampling replicates. **A:** ROHs distributions from the small population; ROHs were called with PLINK. **B:** ROHs distributions from the small population; ROHs were called with RZooRoH. **C:** ROHs distributions from the large population; ROHs were called with PLINK. **D:** ROHs distributions from the large population; ROHs were called with RZooRoH.

### Cattle simulations

Figure 5 shows the comparison of *F*_*ROH*_ estimates, ROHs distributions and fractions of genome correctly assigned within and outside ROHs between WGS and both SNP arrays for the simulated cattle population using *PLINK* and *RZooRoH*. Results are consistent with those obtained with the previous analyses. *F*_*ROH ARRAY*_ are close to *F*_*ROH WGS*_ for both arrays using *PLINK* (points aligned to the equality line) (FIG 5A) and slightly underestimated using *RZooRoH* (points below the equality line) (FIG 5D). Ranks of individuals were always conserved with both arrays (FIG 5A,D; FIG S27). ROHs distributions were biased using both ROHs calling techniques and arrays. With *PLINK*, the mean total length was underestimated for small ROHs (barplots below the horizontal line) and overestimated (barplots above the horizontal line) for large ROHs (FIG 5C) while only the mean total length of small ROHs was underestimated with *RZooRoH* (FIG 5F). As before, the bias was stronger with the small array for both ROHs calling techniques (FIG 5 panels C and F). Approximately 94% of the genome was correctly assigned within or outside ROHs with *PLINK* using both arrays (FIG 5B). On the other hand, around 84% of the genome was correctly assigned within or outside ROHs with *RZooRoH* and the small array and this proportion reached 90% with the large array (FIG 5E).

**Figure 5:**
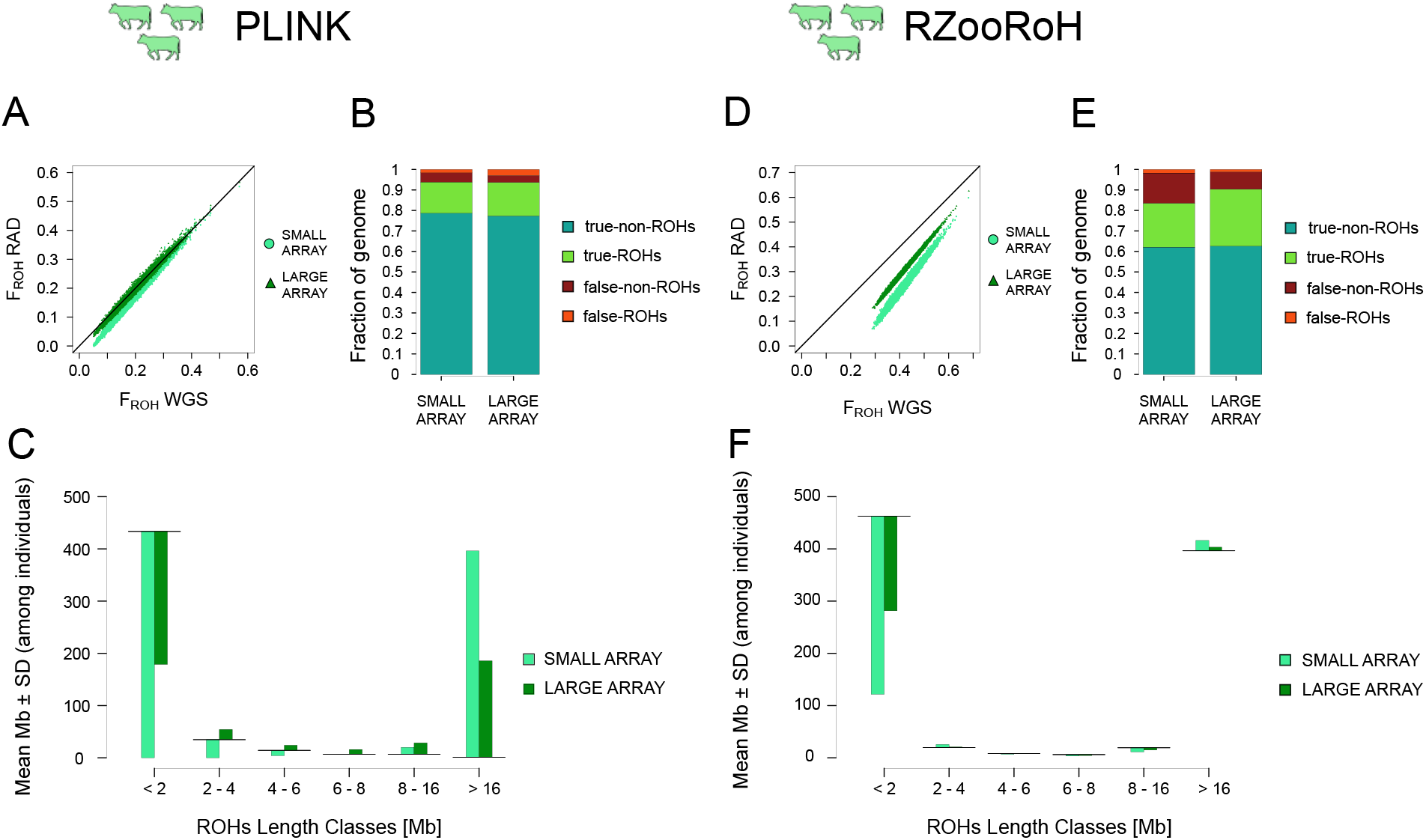
Comparison of ROHs detection between WGS and both SNPs arrays in the cattle population and for both ROHs calling methods. **A:** Comparison of F_ROH_ estimation between WGS and both arrays, ROHs were called with PLINK. **B:** Fraction of genome correctly assigned outside ROHs (true-non-ROHs), correctly assigned within ROHs (true-ROHs), incorrectly assigned outside ROHs (false-non-ROHs) and incorrectly assigned within ROHs (false-ROHs) with both arrays. ROHs were called with PLINK **C:** Comparison of ROHs distributions between WGS and both arrays, ROHs were called with PLINK. **D:** Comparison of F_ROH_ estimation between WGS and both arrays, ROHs were called with RZooRoH. **E:** Fraction of genome correctly assigned outside ROHs (true-non-ROHs), correctly assigned within ROHs (true-ROHs), incorrectly assigned outside ROHs (false-non-ROHs) and incorrectly assigned within ROHs (false-ROHs) with both arrays. ROHs were called with RZooRoH **F:** Comparison of ROHs distributions between WGS and both arrays, ROHs were called with RZooRoH.

## DISCUSSION

### Summary

We investigated whether reduced representation of WGS such as RAD-sequencing and SNPs arrays are reliable for ROHs analyses using either the observational-based approach implemented in *PLINK* (Chang et al., 2015; Purcell et al., 2007) or the model-based approach implemented in *RZooRoH* (Bertrand et al., 2019; Druet & Gautier, 2017). We show that reduced genomic representations possibly lead to underestimated individuals *F*_*ROH*_ and to biased ROHs distributions. Despite this, the rank of individual *F*_*ROH*_ was conserved when *F*_*ROH REDUCED*_ was estimated with a sufficient density of SNPs. The model-based approach required a smaller proportion of the genome to be sequenced to obtain reliable estimates of individual *F*_*ROH*_. In addition, we observed a biased estimation of ROH length distribution, independently of the reduced representation used with *PLINK*. On the contrary, we observed this bias only with RAD-sequencing and, to a lesser extent, with the small array with *RZooRoH*. Finally, we show in supplementary material that *F*_*HOM*_, a non-ROH based estimate of individual *F*, is as accurate as *F*_*ROH*_ estimated with the model-based approach for the same SNP density.

### ROHs estimates with WGS

First, we want to briefly comment on the two methods we used for ROHs calling. With WGS, both softwares yielded different results. We showed that *PLINK* mostly identifies IBD segments coalescing less than 100 (overlapping) generations ago while *RZooRoH* captures more ancient inbreeding events (FIG S2, S3 & S4). This is expected as *PLINK* uses a minimum length threshold for a homozygous segment to be classified as a ROH which only allows to detect recent inbreeding. From the formula presented in Thompson (2013): l 100/2g with l as the length of the IDB segment in centimorgan (cM) and g as the time in generation back to the coalescence event, our threshold corresponds to 0.1 cM (100KB) in humans and thus inbreeding events 500 reproduction events ago on average (with a very large variance though as shown with simulations by (Speed & Balding (2015)). With *RZooRoH*, we did not set any size threshold and chose HBD classes which allowed the software to capture older inbreeding events. It is important to remember that even with WGS, ROHs detection is still challenging. Indeed, beside the ROHs calling method, many parameters can influence the result of ROHs calling. For instance, the use of a minimum size threshold can be chosen to ensure that the homozygous regions detected are truly IBD and not Identical-by-State (IBS). This choice often relies on the hypothesis that small segments come from ancient inbreeding events. However, it has been shown with simulations that small segments can also result from recent inbreeding events (Speed & Balding, 2015), suggesting that neglecting smaller regions might lead to an underestimation of the inbreeding status of the individual or the population. Another important parameter is whether mutations (and sequencing errors) should be considered by allowing heterozygous SNPs in IBD segments. Even though it will only affect the sum of IBD segments trivially, how many heterozygous markers are to be allowed can greatly differ among studies. To summarize, no consensus exist nowadays on which method and parameters are the best and further investigation is needed.

### Estimation of *F*_*ROH*_

Reduced genomic representations resulted in consistent relative estimates of *F*_*ROH*_ providing a large enough proportion of the genome is sequenced but unreliable absolute values. With both arrays and when a large portion of the genome is sequenced with RAD-sequencing, ranks of inbreeding were conserved among individuals. Our results are consistent with Kardos et al. (2018) who showed (see this paper supplementary material) that *F*_*ROH*_ estimated with 10,000 loci, and a home-made script based on a likelihood ratio method (adapted from Pemberton et al., 2012), are similar to *F*_*ROH*_ estimated with WGS in an inbred wild wolf population. Duntsch et al. (2021) compared ROHs estimates from *PLINK* and *RZooRoH* with RAD-sequencing, a custom-made array and WGS in few hihi (*Notiomystis cincta;* a non-model bird species) individuals. They found conserved individual inbreeding ranks with reduced representation. This suggest that reduced genomic representation can be used to estimate inbreeding from ROHs, if the aim is to compare individuals within or between populations.

With low marker densities *F*_*ROH*_ estimates were often underestimated. The portion of inbreeding which is missing in that case is not random: it is ancient inbreeding. Indeed, recent inbreeding, represented by long ROHs segments is easier to capture; with lower marker density you are no longer able to capture small segments (Druet & Gautier, 2017). Consequently, the base population moves towards more recent ancestors. This is supported by the study from Sole et al. (2017) where the authors compared SNPs arrays of different sizes in cattle and showed that all arrays capture the same levels of recent inbreeding but higher densities allow to capture more ancient inbreeding.

Inbreeding coefficients are often used for inbreeding depression studies, which are key for understanding the evolution of populations and for conservation managements in endangered species (Lynch & Walsh, 1998). If the rank of individuals’ *F* is conserved, it ensures that the direction of the correlation between inbreeding estimates and phenotypes (the sign of the regression slope) and thus the general effect of inbreeding on the trait is correctly estimated. However, underestimating the overall inbreeding coefficients can lead to an underestimation of the magnitude of inbreeding depression in a population (the regression slope absolute value). We showed in supplementary material that another inbreeding coefficient: *F*_*HOM*_ performed as well as model-based *RZooRoH* approach with similar SNPs densities. This last result is consistent with another study which showed that 5,000 markers are sufficient to obtain genomic kinship values similar to pedigree estimates (Goudet et al., 2018). Hence, we strongly advise to use SNP independent measures (i.e. not based on IBD segments) or model-based ROHs approaches for inbreeding studies when SNP density is low (see below). Another solution would be to use imputation of non-sequenced genomic regions to increase the SNP density but this implies the availability of a reference panel not easily accessible for non-model species (Marino et al., 2021).

### ROHs distributions

ROHs distributions are commonly used to describe the demographic history of the population (Bosse et al., 2012; Ceballos et al., 2018; Kirin et al., 2010). Indeed, Thompson (2013) proposed a formula to link the length of an IBD segment to the number of generations back to the common ancestor (and thus the inbreeding event in the case of a ROH). Our study suggests that with most reduced genomic representations, ROHs lengths values should not be trusted, especially in small populations, meaning that age estimation for these IBD segments is currently impossible. However, as we could expect this bias to be similar among the different populations, relative comparisons between populations from a given study or genotyped with the same sequencing method lead to reliable results as performed by Kirin et al. (2010). In the latter, the authors used the HGDP dataset genotypes (from 2009) with Illumina 650Y product and compared ROHs distributions among various humans’ populations from all around the world. The populations which underwent recent inbreeding events such as West and South Asians and Oceanians harbored higher fraction of genome within long ROHs, consistent with what we know about human populations’ history. Similarly, Mastrangelo et al. (2016) compared ROHs distributions from three dairy cattle breeds genotyped with the ‘small array’ (Illumina BovineSNP 50 beadchip) in order to compare the different breeds and to assess their inbreeding status. The authors found that Italian Holstein individuals harbored high number of short ROHs suggesting that inbreeding in this breed is mostly caused by ancient relatedness within the population rather than recent mating between relatives. On the contrary, individuals from two other local breeds: Modicana and Cinisara harbored high number of large ROHs, suggesting the presence of recent mating events between close relatives. The authors concluded that the implementation of a closely monitored breeding program aiming at reduced consanguinity was necessary in these two local breeds.

We observed an underestimation of short and overestimation of long ROHs with reduced genomic representations. We hypothesized that reduced representations techniques (or reduced number of SNPs) tend to merge small adjacent ROHs into larger ones. Our results are consistent with Duntsch et al. (2021) who suggested that very long homozygous regions identified with both a custom array and RAD-sequencing in hihi (*Notiomystis cincta*) are expected to be artefacts of smaller merged IBD regions. The authors concluded that higher density of markers is required to accurately identify homozygous stretches. This result is also confirmed by the correct assignment of the genome within and outside ROHs that we obtained: close to 90% with the small and to 95% with the large array. The remaining incorrectly assigned regions correspond mostly to either very small sparse ROHs (much harder to detect with small SNPs densities (Druet & Gautier, 2017; Kardos et al., 2015; Sole et al., 2017)) or segments in between adjacent ROHs. This trend indicates that IBD regions are overall correctly detected with both SNPs arrays and thus suggest that they can be confidently used for homozygosity mapping studies.

### PLINK *vs* RZooRoH

With reduced genomic representations, accurate results of ROHs calling depends on the fraction of genome sequenced, which drastically differ between both ROHs calling softwares. *RZooRoH* required a fraction of the genome 10 times smaller compared to *PLINK* for accurate ROHs detection. It is expected that model-based approaches perform better with low SNPs-densities as the distances between SNPs are taken into account in the model (Bertrand et al., 2019). Hence, we strongly recommend *RZooRoH* when working with reduced genomic data, even though it requires more computational resources.

Interestingly, *RZooRoH* resulted in more false-ROHs (i.e. false positive) compared to *PLINK*, in the small population and with the small and to a lesser extend the large arrays. Since we observed an underestimation of ROHs smaller than 2MB but an overestimation of ROHs between 2MB and 6MB for the small array, we hypothesize than this is the results of the merging of small adjacent ROHs we mentioned in the previous paragraph. Regarding the large array, we observe a slight overestimation of all ROHs length classes with *RZooRoH* and thus we believe that this increase of false-ROHs is due both to merging of small adjacent ROHs and spurious identification of small false-ROHs in non IBD regions.

In addition, in the large population and at similar densities (i.e. both arrays) *PLINK* yielded a higher fraction of genome correctly assigned compared to *RZooRoH*. In this population, fractions of genome incorrectly assigned, were mostly false-non-ROHs (i.e. false-negative) using *RZooRoH*. From the ROHs distributions, we know that the ROHs segments which are missing are mostly short ROHs smaller than 2MB. Since *RZooRoH* does not impose any minimum size threshold (while we did with *PLINK*), these missing ROHs segments are likely to be smaller than those detected with *PLINK*. Consequently, it is expected that fraction of false-non-ROHs is higher with *RZooRoH* compared to *PLINK* as short fragments are harder to detect and require higher densities (Druet & Gautier, 2017; Kardos et al., 2015; Sole et al., 2017). Finally, we want to stress that fine-tuning of the number and rates of inbreeding classes when constructing the *RZooRoH* model can help to better detect specific lengths classes ROHs.

### Effect of population size

Not only did the ROHs calling method influence the minimum proportion of genome required, but population size also played a role. The larger population required a smaller fraction of genome to obtain accurate ROHs calling. In our simulations, the larger population harbored a higher genetic variation and thus a higher number of SNPs. Consequently, we were more likely to have SNPs in the subsampled regions with both RAD-sequencing and SNPs arrays. Since the simulations scheme forced mating between related individuals and, since we performed a random stratified individual sampling independently of the effective size of the population, we do not expect ROHs distributions to actually reflect both populations “true” ROHs distributions. Indeed, within the subsampled individuals, the large population harbors as much long ROHs as the small population which itself harbors higher numbers of small ROHs compared to the large population. The latter is due to the smaller effective size and thus the higher coancestry between individuals in the small population. Consequently, in this study we cannot quantify the true effect of population size (other than the number of SNPs) on ROHs detection.

### Limitations

Even though we made efforts to perform realistic simulations, additional studies with non-simulated data such as performed by Duntsch et al. (2021) in hihi will be important to strengthen our findings. In addition, we used the softwares recommended default settings for ROHs calling. We note that previous studies (Duntsch et al., 2021; Meyermans et al., 2020; Mueller et al., 2022) showed that changing these settings could lead to different results. For instance, Mueller et al., (2022) showed adapting parameters might allow to get similar results between WGS and reduced genomic representations with smaller fractions of genomes: the authors used *PLINK* to call ROHs with RAD-sequencing. They then tested ROH calling with several different settings for three individuals for which they also had WGS data and extracted the settings which best conserved the rank of inbreeding found with WGS. However, varying settings need to be done with caution, especially with *PLINK*. Indeed, different settings can increase the number of ROHs detected but also the likelihood of non-correct calls and maybe bias the individuals inbreeding ranking (Meyermans et al., 2020). We want to stress that the fractions of the genome sequenced presented in the present study are indicative and do not correspond to “true” proportion of genome sequenced needed to obtain meaningful results. Indeed, they come from ideal simulated data and this fraction will be lower after quality filtering with non-simulated data. Finally, in this study we only focused on inbreeding coefficients and ROHs distributions estimation; we did not investigate whether variants types (i.e. synonymous or non-synonymous mutations) are similarly enriched in particular ROHs length categories between WGS and reduced genomic representations.

## Conclusion

Using simulated data, we compared *F*_*ROH*_ estimates and ROHs distributions between WGS and reduced genomic representations with large sample sizes and our results are concordant with previous studies using empirical data. ROHs based individual inbreeding coefficients can be correctly estimated with reduced genomic representations when the SNP density is above 20 SNPs per Mb with *PLINK* and above 3 SNPs per Mb with *RZooRoH*. We found that *F*_*HOM*_, a genomic estimate of inbreeding coefficients not based on ROHs, is as accurate as the model-based ROHs estimate, for a fraction of the computing time. We would therefore recommend using independent SNPs-based genomic estimates such as *F*_*HOM*_ for inbreeding quantification with reduced genomic representation, unless the number of individuals analyzed is too small to allow a correct estimation of the population alleles frequency or mean coancestry. Regarding ROHs distributions, even though the majority of the genome is correctly assigned within and outside ROHs, the lengths of the homozygous segments are biased with a deficit of short and an excess of long ROHs. This bias was observed for all reduced genomic data for *PLINK* and with RAD-sequencing for *RZooRoH*. With the low-density array (i.e. small array) only ROHs smaller than 2MB were largely underestimated in the large population with *RZooRoH*. This can still allow comparing populations analyzed with the same methodology but prevent comparing ROH distributions from studies using different reduced genomic methods. To conclude, we find little advantages in using ROHs based measures of inbreeding with low SNPs densities and reduced representations and show that only model-based approaches with high SNPs densities can be used for ROHs distributions quantification.

## ACKOWLEDGMENTS

We are grateful to the Association Wallonne de l’Elevage (awé) for accepting to share their cattle pedigree with us and to Tom Druet who kindly asked them on our behalf and provided feedbacks on this manuscript. This study was funded by the Swiss National Science Foundation with grant 31003A_179358 to JG. Open Access Funding provided by University of Lausanne.

## DATA ACCESSIBILITY

All scripts necessary to generate genomic data and perform analyses used in this study are available on GitHub: https://github.com/EleonoreLavanchy/ROHsReducedRep.

## CONFLICT OF INTEREST STATEMENT

The authors declare no conflict of interest.

## AUTHOR CONTRIBUTIONS

EL and JG conceptualized the study. EL conducted the study, performed all simulations and analyses and drafted the manuscript. EL and JG wrote and revised the manuscript.

## Notes

### Competing Interest Statement

The authors have declared no competing interest.

