## Supplementary Material for "Effect of reduced genomic representation on using runs of homozygosity for inbreeding characterization"

**SUPPLEMENTARY MATERIAL & METHODS**

**Simulations: small and large populations**

The genetic map was simulated with *FREGENE* (Chadeau-Hyam et al., 2008) with default parameters to mimic a human-like recombination map. Input files with default parameters can be accessed while downloading the *FREGENE* software in *Fregene/Example/data/*. Only the *<CHROMO_LENGTH>* parameter was modified to simulate chromosomes with size of 100KB.

We performed two rounds of simulations with *SLiM3* (Haller & Messer, 2019) to force all chromosomes from the same simulation replicate to have the same history/pedigree. For each simulation replicate, the aim of the first round was to generate the pedigree which was then “applied” 30 times (for 30 chromosomes) in the second round of simulations. In the first round of simulations, we used a non-Wright-Fisher model with overlapping generations: individuals ages varied between 0 and 3 years old. Ages were uniformly assigned to individuals at the first generation. The population size was regulated at each generation at the end of the simulation cycle: only *Ne* individuals (i.e. 1,000 for the small and 10,000 for the large population) with the higher relative fitness survived through the next generation. In addition to the density-dependency fitness, individual fitness was relative to age: probability to survive to the next generation was 0.9 for 0-year old individuals, 0.7 for 1-year old individuals, 0.5 for 2-years old individuals and 0 for 3-years old individuals. A reproduction callback (which defines reproduction events in SLiM) was called at each generation: *Ne* individuals (i.e. 1,000 for the small and 10,000 for the large population) were sampled for reproduction (without replacement). Individual probabilities of being selected as a mate were age-dependent: 0 for 0 years old individuals, 0 for 1-year old individuals, 0.6 for 2-years old individuals and 0.4 for 3-years old individuals. For each of these *Ne* individuals, a mate was then selected among the other adults (i.e. older than 1-year old) of the population, thus creating one new individual in the population. For each potential mate, the probability of being chosen as a mate was equal to 10 times the relatedness (estimated with *SLiM*) between the potential mate and the focal individual + 0.01 multiplied by 0.6 for 2-years old individuals and 0.4 for 3-years old individuals. The first part was to ensure that some inbreeding occurred at each generation. Mating and death events were recorded for generation. During the second round of simulations, these mating and deaths events were simply applied to the 30 chromosomes (each with a different genetic map).

The burn-in were performed with *Python* via *recapitation* in *msprime* (Kelleher et al., 2016), with a constant recombination rate of 1e^-8^. Mutations were added at the end of the simulation based on a human-like mutation rate of ${2.5}^{e-8}$.

**Simulations: cattle population**

The genetic map from male Holstein cattle was obtained from a study from Qanbari & Wittenburg (2020). Chromosome sizes correspond to the true chromosome sizes from *Bos Taurus*.

The history of the simulation was inspired by a paper from Frantz et al. (2020). For this population, we first simulated the burn-in with *msprime:* We simulated two populations: a wild population with 50,000 individuals and a domestic population with 1,500 individuals which split 10,000 generations ago. The migration rate between both populations was constant and equal to 3e^-5^. During this burn-in, the recombination rate was constant and equal to 1e^-8^. At the end of the burn-in, 200 individuals were randomly sampled to mimic the strong artificial pressure applied 200 generations ago during many current breeds creation. Individual metadata were modified to fit a non-Wright-Fisher model: half of the individual were assigned a female gender and the other half a male gender. Individual ages were randomly assigned between 0 and 3 years old.

The last 200 generations were performed in *SLiM3*. Since our real pedigree only covered 57 years maximum, we performed a first round of simulation and recorded all mating events. We then used this simulated pedigree to complete the real one: each founder of the real pedigree (i.e. each individual from which parents are unknown) was randomly linked to a simulated individual from the same generation, consequently receiving his genealogy (from the simulated pedigree). In other words, we used the simulated genealogies to complete the real ones. Afterwards, we trimmed this completed pedigree to keep only the individuals and mating events resulting in the individuals from the last generation of the real pedigree. Then, we ran a second round of simulations were we simply applied this new complete pedigree to all chromosome of the cattle genome in *SLiM*. Finally, we used *msprime* to add all mutations with a mutation rate of 2.5e^-8^ and to subsample the individuals from the last generation only. This corresponds to our WGS data for the cattle population.

**SNPs-independent measures of inbreeding: F_HOM_**

To test the performance of a SNPs-independent based inbreeding coefficient, we estimated F_HOM_, implemented in the *--het* method from *PLINK* with both WGS and at low SNPs-densities.

**Heterozygosity along the genome**

To visualize the effect of the sequencing method on ROHs detection, we plotted the heterozygosity along the genome as proposed by Kardos et al. (2018) (Kardos et al., 2018) with each ROHs detected for specific genomic regions in one replicate of the subsampling and simulation replicates. Heterozygosity was calculated as proportion of heterozygous sites per individual in 50Kb (20Kb-overlapping) sliding windows.

**SNP Density**

We calculated SNP densities with *VCFtools* (Danecek et al., 2011) method: *--SNPdensity* as the number of SNPs per each windows of 1Mb. We then estimated the mean SNP density of each replicate as the mean density among the windows.

**SUPPLEMENTARY RESULTS**

**Figure S1**

**Figure S1**: Comparison of F_PED_ estimated with the last 15 generations , F_PLINK 100KB_ and F_RZooRoH_ in the three simulated populations.

Figure S1 shows the relationship between the three inbreeding coefficients: : F_PED_, F_ROH PLINK_ and F_ROH RZooRoH_ in the three populations. In both the small and large populations, the three measures of inbreeding: F_PED_, F_ROH PLINK_ and F_ROH RZooRoH_ yield similar results of individual inbreeding quantification: even though F_ROH RZooRoH_ always detect more inbreeding compared to *PLINK* and the pedigree approach, the rank of inbreeding are mostly conserved between the three approaches. In the cattle population however, F_PED_ gives very different results compared to F_ROH PLINK_ and F_ROH RZooRoH_. This is probably because inbreeding is mostly due to ancient relatedness in this population: recent mating events are not between closely related individuals and the pedigree was estimated with the last 15 generations and thus cannot detect any inbreeding events older than that.

**Figure S2**

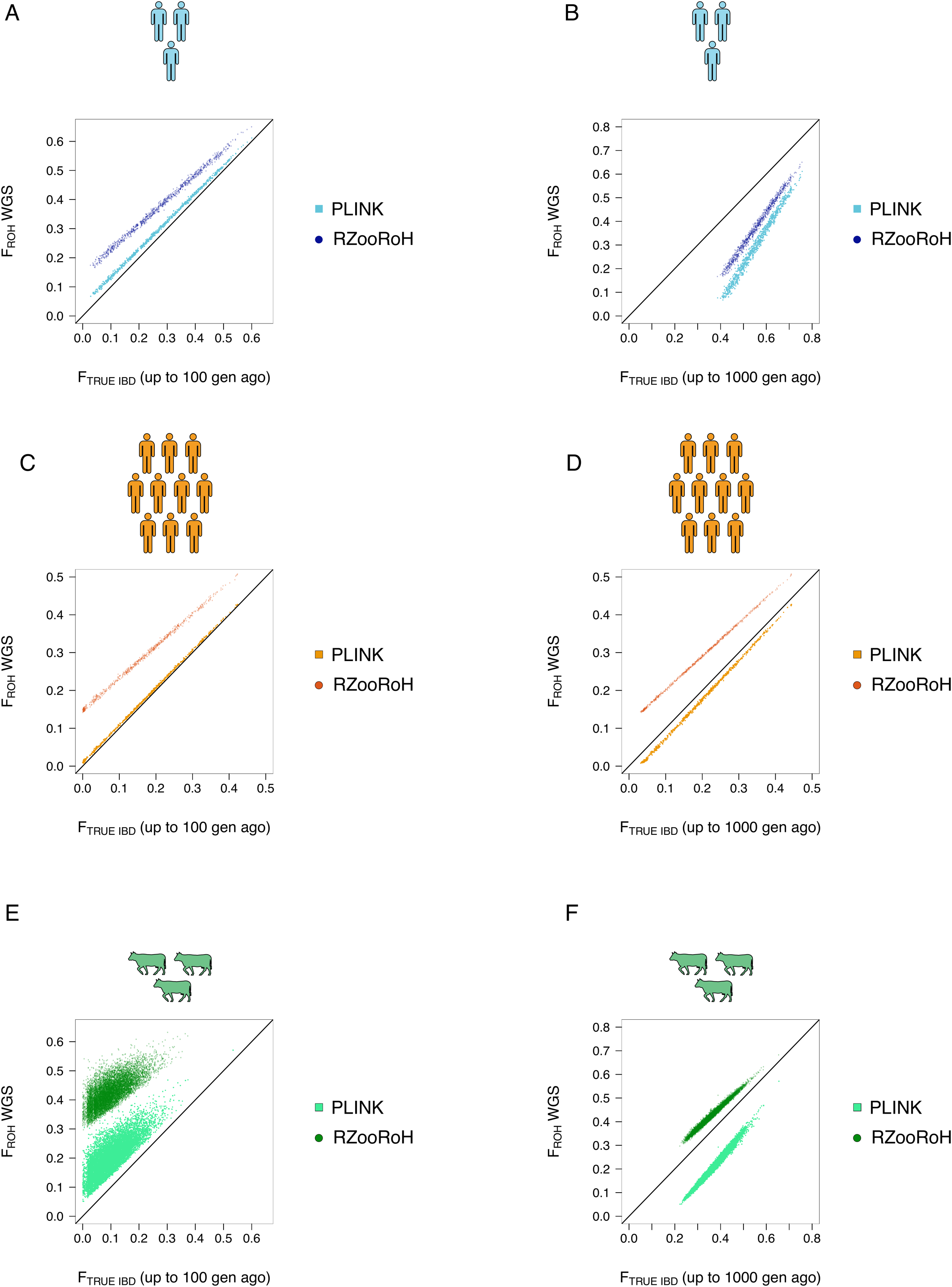

**Figure S2:** Comparison between F_ROH_ estimated with WGS data and proportion of genome within true IBD segments. **A:** Small population, a genomic segment was considered as IBD if both haplotype within the individuals coalesced less than 100 generations ago. **B:** Small population, a genomic segment was considered as IBD if both haplotype within the individuals coalesced less than 1000 generations ago. **C:** Large population, a genomic segment was considered as IBD if both haplotype within the individuals coalesced less than 100 generations ago. **D:** Large population, a genomic segment was considered as IBD if both haplotype within the individuals coalesced less than 1000 generations ago. **E:** Cattle population, a genomic segment was considered as IBD if both haplotype within the individuals coalesced less than 100 generations ago. **F:** Cattle population, a genomic segment was considered as IBD if both haplotype within the individuals coalesced less than 1000 generations ago.

In Figure S2, F_ROH_ estimated with both *PLINK* and *RZooRoH* are compared to the true fraction of genome within IBD segments in the three populations. A segment is considered IBD if both copies come from the same ancestor within the last 100 or 1,000 generations. In the small and large populations, F_ROH PLINK_ is very close to the true inbreeding coefficient with IBD segments coming from 100 generation ago or younger. In the small population, the difference between the true fraction of the genome within IBD segments from 100 generations ago and true fraction of the genome within IBD segments 1,000 generations ago is smaller compared to what we observe in the large population. We believe it is because in the large population, there is more ancient inbreeding compared to the small population. In the small population, both *PLINK* and *RZooRoH* underestimate the fraction of genome which is IBD coming from up to 1,000 generations ago. In the large population, RZooRoH largely overestimates the fraction of genome within IBD segments up to 1,000 generations ago. It could be that *RZooRoH* detects IBD segments from which the common ancestor is older than 1,000 generations ago. However we’ll see in Figure S4 that fraction of genome incorrectly assigned within ROHs is higher for *RZooRoH* in the large population, suggesting that it is mostly due to false-positive: i.e. some ROHs which have been detected are actually not IBD segments. Finally, in the cattle population, we can see that both F_ROH PLINK_ and F_ROH RZooRoH_ show low correlation with the fraction of genome within IBD segments from which the common ancestor was younger or equal to 100 generations ago. On the contrary, the correlations are higher with fraction of genome within IBD segments from which the common ancestor was younger or equal to 1,000 generations ago. This confirms what we mentioned before: in the cattle population inbreeding is principally due to ancient relatedness among individuals: there are very few recent inbreeding events (i.e. mating between individuals who share a recent common ancestor).

**Figure S3**

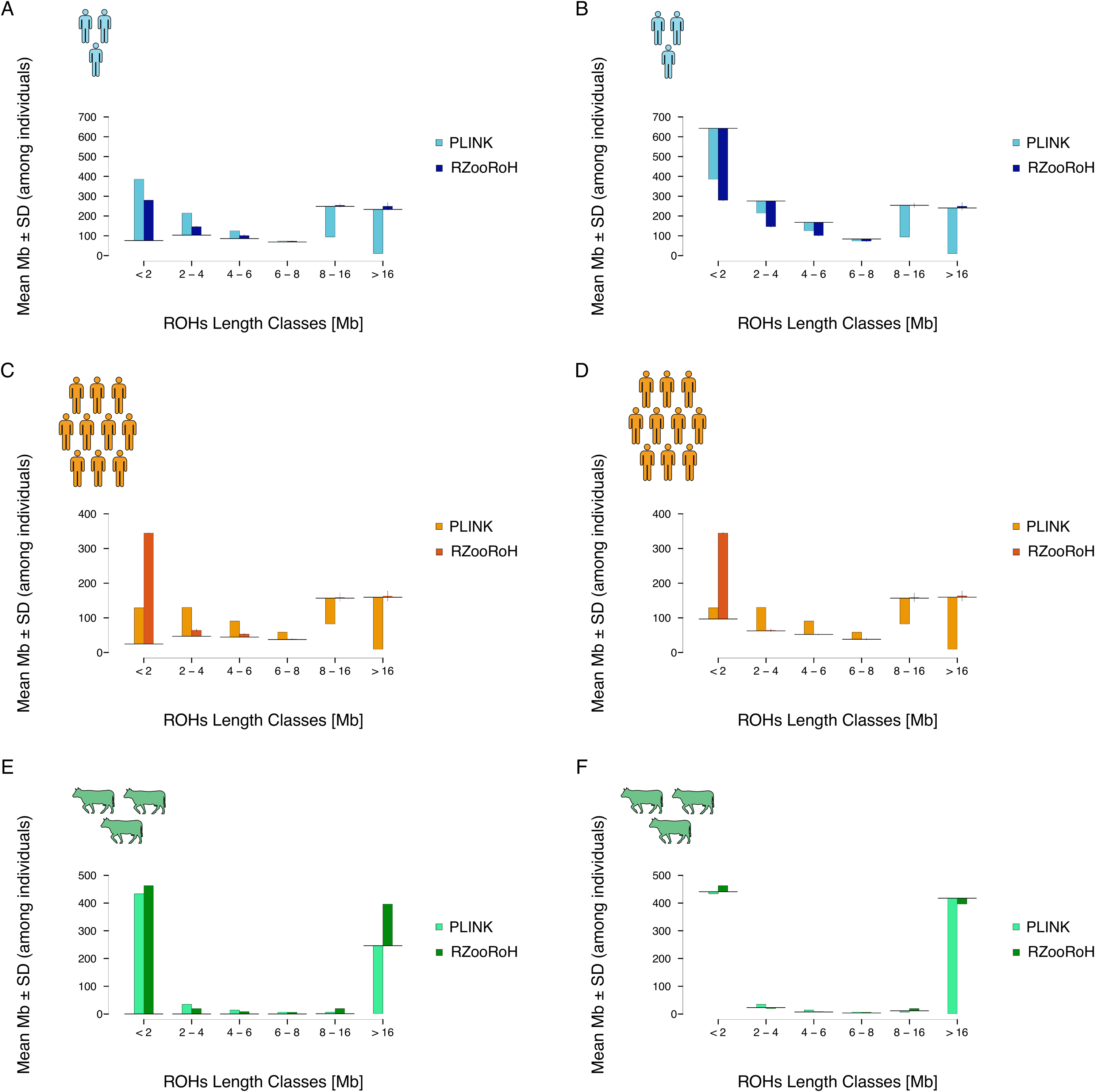

**Figure S3:** Comparison between ROHs distributions estimated with WGS data and true IBD segments distributions. **A:** Small population, a genomic segment was considered as IBD if both haplotype within the individuals coalesced less than 100 generations ago. **B:** Small population, a genomic segment was considered as IBD if both haplotype within the individuals coalesced less than 1000 generations ago. **C:** Large population, a genomic segment was considered as IBD if both haplotype within the individuals coalesced less than 100 generations ago. **D:** Large population, a genomic segment was considered as IBD if both haplotype within the individuals coalesced less than 1000 generations ago. **E:** Cattle population, a genomic segment was considered as IBD if both haplotype within the individuals coalesced less than 100 generations ago. **F:** Cattle population, a genomic segment was considered as IBD if both haplotype within the individuals coalesced less than 1000 generations ago.

Figure S3 shows the ROHs segments distributions compared to the distribution of true IBD segments (estimated as before: with the common ancestor younger or equal to 100 or 1,000 generations ago). Concerning IBD segments for which the common ancestor is younger than 100 generations ago, PLINK overestimates small ROHs total lengths and underestimates large ROHs total lengths in both the small and large populations. On the contrary, RZooRoH correctly estimates all total length except for ROHs smaller than 2MB for which it overestimates the total length. Concerning IBD segments for which the common ancestor is younger than 1,000 generations ago, *PLINK* underestimates the ROHs total lengths in all ROHs length classes in the small population and overestimates small ROHs but underestimates large ROHs total lengths in the large population. In contrast, *RZooRoH* only underestimates the total length for ROHs smaller than 6MB but correctly estimate the total length for large ROHs in the small population. In the large population only ROHs smaller than 2MB are overestimated with *RZooRoH*. Finally, concerning the cattle population, true IBD segments for which the common ancestor is younger than 100 generations ago only fall into large segments. *PLINK* and *RZooRoH* both largely overestimates the total length of small ROHs (<2MB) but *PLINK* underestimates the total length of large ROHs (>16MB) while they are overestimated by *RZooRoH*. Concerning true IBD segments for which the common ancestor is younger than 1,000 generations ago, RZooRoH shows accurate ROHs distributions while PLINK only underestimates large ROHs (> 16Mb).

**Figure S4**

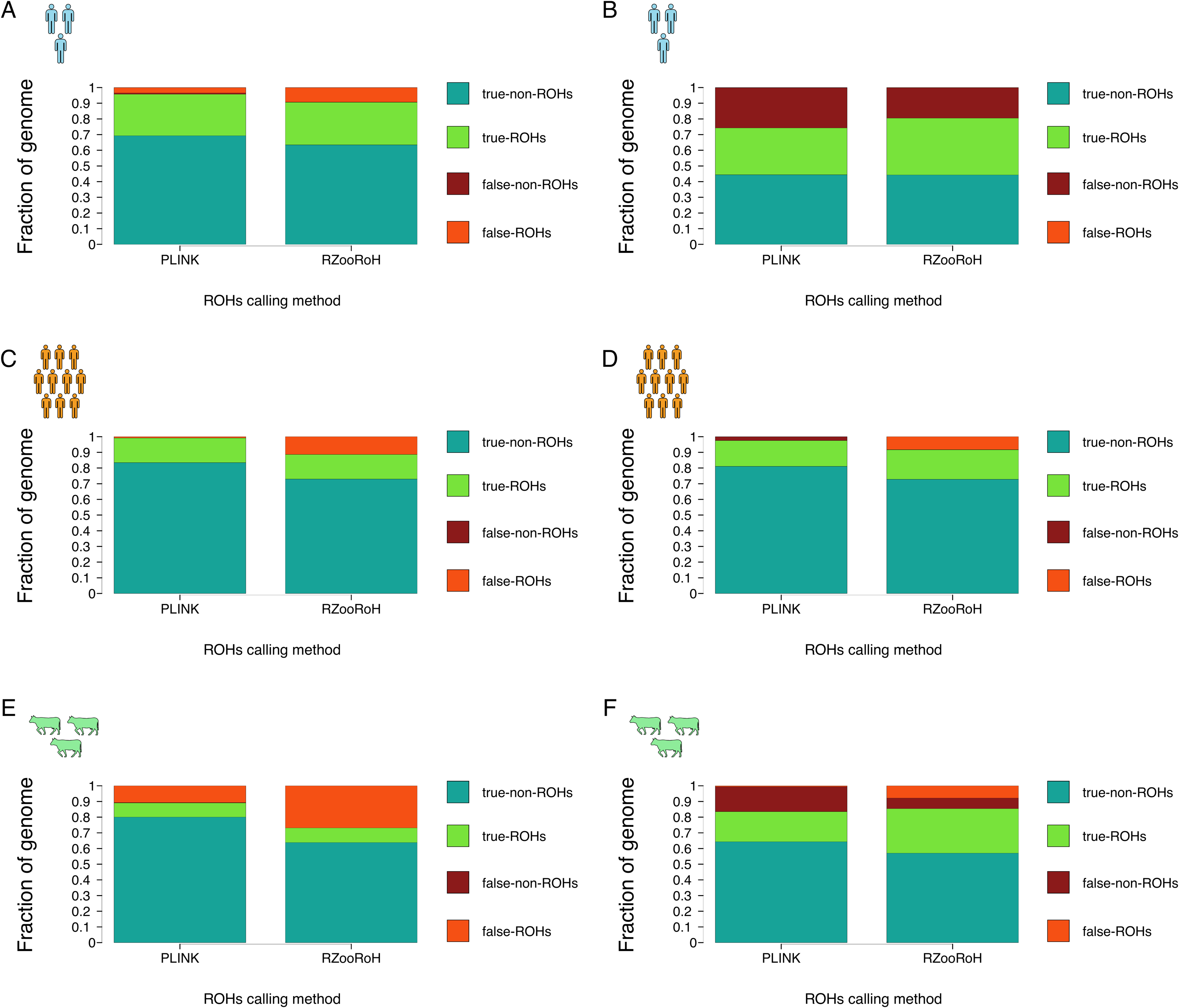

**Figure S4:** Fraction of genome correctly assigned outside ROHs (true-non-ROHs), correctly assigned within ROHs (true-ROHs), incorrectly assigned outside ROHs (false-non-ROHs) and incorrectly assigned within ROHs (false-ROHs). **A:** Small population, a genomic segment was considered as IBD if both haplotype within the individuals coalesced less than 100 generations ago. **B:** Small population, a genomic segment was considered as IBD if both haplotype within the individuals coalesced less than 1000 generations ago. **C:** Large population, a genomic segment was considered as IBD if both haplotype within the individuals coalesced less than 100 generations ago. **D:** Large population, a genomic segment was considered as IBD if both haplotype within the individuals coalesced less than 1000 generations ago. **E:** Cattle population, a genomic segment was considered as IBD if both haplotype within the individuals coalesced less than 100 generations ago. **F:** Cattle population, a genomic segment was considered as IBD if both haplotype within the individuals coalesced less than 1000 generations ago.

Figure S4 shows the fraction of genome correctly and incorrectly assigned within and outside the “true” IBD segments for the three populations. Concerning IBD segments for which the common ancestor is younger or equal to 100 generations ago, more than 95% and 90% of the genome is correctly assigned with *PLINK* and *RZooRoH* respectively in both populations. For *PLINK*, the remaining 5% are half false-ROHs and half false-non-ROHs. False-ROHs are either small non-IBD regions in between adjacent IBD segments that were falsely incorporated in the ROH or Identical-by-state (IBS but not IBD) segments. False-non-ROHs can be small fragments which were not detected either due to small SNPs density in the corresponding regions or a size smaller than 100MB. Concerning *RZooRoH*, genome incorrectly assigned within or outside true IBD segments is mostly composed of false-ROHs. These are probably IBS (but not IBD) segments which coalesce more than 100 generations ago. Concerning IBD segments for which the common ancestor is younger or equal to 1,000 generations ago, fraction of genome correctly assigned within and outside ROHs are lower for both methods and populations. In the small population, both *PLINK* and *RZooRoH* results in a lot of false-non-ROHs, i.e. IBD segments were not detected as a ROH. This is probably because old IBD fragments can be in monomorphic regions where the SNP density is thus very low or equal to zero or because these old IBD segments are small and thus undetected (it is the case for all IBD segments smaller than 100KB for *PLINK*). Concerning the large population, we observe the same pattern as before: *PLINK* assigns most of the genome correctly with few false-non-ROHs and almost no false-ROHs while *RZooRoH* results in higher numbers of false-ROHs. For PLINK, the difference compared to the small population is probably because there are fewer monomorphic genomic regions in the large population compared to the small population and thus higher SNP densities which lead to better ROH detection. Concerning RZooRoH, as before we expect that most of these segments are IBS rather than IBD and were thus falsely labeled as ROHs. Finally, concerning the cattle population, both methods resulted in high fraction of genome in false-ROHs for IBD segments younger than 100 generations. As mentioned before, this is probably because inbreeding is mostly ancient in this population. Consequently, many ROHs are IBD fragments older than these 100 generations. For IBD segments for which the common ancestor was younger or equal to 1,000 generations, the fraction of genome incorrectly assigned with *PLINK* is mostly false-non-ROHs, i.e. probably IBD segments smaller than 100KB. On the contrary, fraction of genome incorrectly assigned within or outside ROHs was half false-ROHs and half false-non-ROHs for *RZooRoH*.

**Figure S5**

**Figure S5:** Correlation between F_ROH_ estimated with the reduced dataset and F_ROH_ estimated with WGS data according to SNP density in the reduced dataset for both the small and large population. **A**: ROHs have been called with PLINK; **B:** ROHs have been called with RZooRoH.

**Figure S6**

**Figure S6:** Distributions of the difference in inbreeding rank between WGS and RAD-sequencing in the small population with different proportion of genome sequenced. ROHs were called with PLINK, minimum size 100KB. Individual inbreeding rank were estimated for the sequencing techniques and the difference between both was obtained for each individual in each subsampling replicate. The distribution of these differences is represented in this figure. **A:** 3% of genome sequenced; **B:** 4% of genome sequenced; **C:**5% of genome sequenced; **D:**15% of genome sequenced.

**Figure S7**

**Figure S7:** Distributions of the difference in inbreeding rank between WGS and RAD-sequencing in the large population with different proportion of genome sequenced. ROHs were called with PLINK, minimum size 100KB. Individual inbreeding rank were estimated for the sequencing techniques and the difference between both was obtained for each individual in each subsampling replicate. The distribution of these differences is represented in this figure. **A:** 0.25% of genome sequenced; **B:** 0.33% of genome sequenced; **C:**0.5% of genome sequenced; **D:**1% of genome sequenced.

**Figure S8**

**Figure S8:** Distributions of the difference in inbreeding rank between WGS and RAD-sequencing in the small population with different proportion of genome sequenced. ROHs were called with RZooRoH, 3 HBD classes. Individual inbreeding rank were estimated for the sequencing techniques and the difference between both was obtained for each individual in each subsampling replicate. The distribution of these differences is represented in this figure. **A:** 0.05% of genome sequenced; **B:** 0.15% of genome sequenced; **C:**0.5% of genome sequenced; **D:**1% of genome sequenced.

**Figure S9**

**Figure S9:** Distributions of the difference in inbreeding rank between WGS and RAD-sequencing in the small population with different proportion of genome sequenced. ROHs were called with RZooRoH, 3 HBD classes. Individual inbreeding rank were estimated for the sequencing techniques and the difference between both was obtained for each individual in each subsampling replicate. The distribution of these differences is represented in this figure. **A:** 0.002% of genome sequenced; **B:** 0.008% of genome sequenced; **C:** 0.042% of genome sequenced; **D:** 1% of genome sequenced.

**Figure S10**

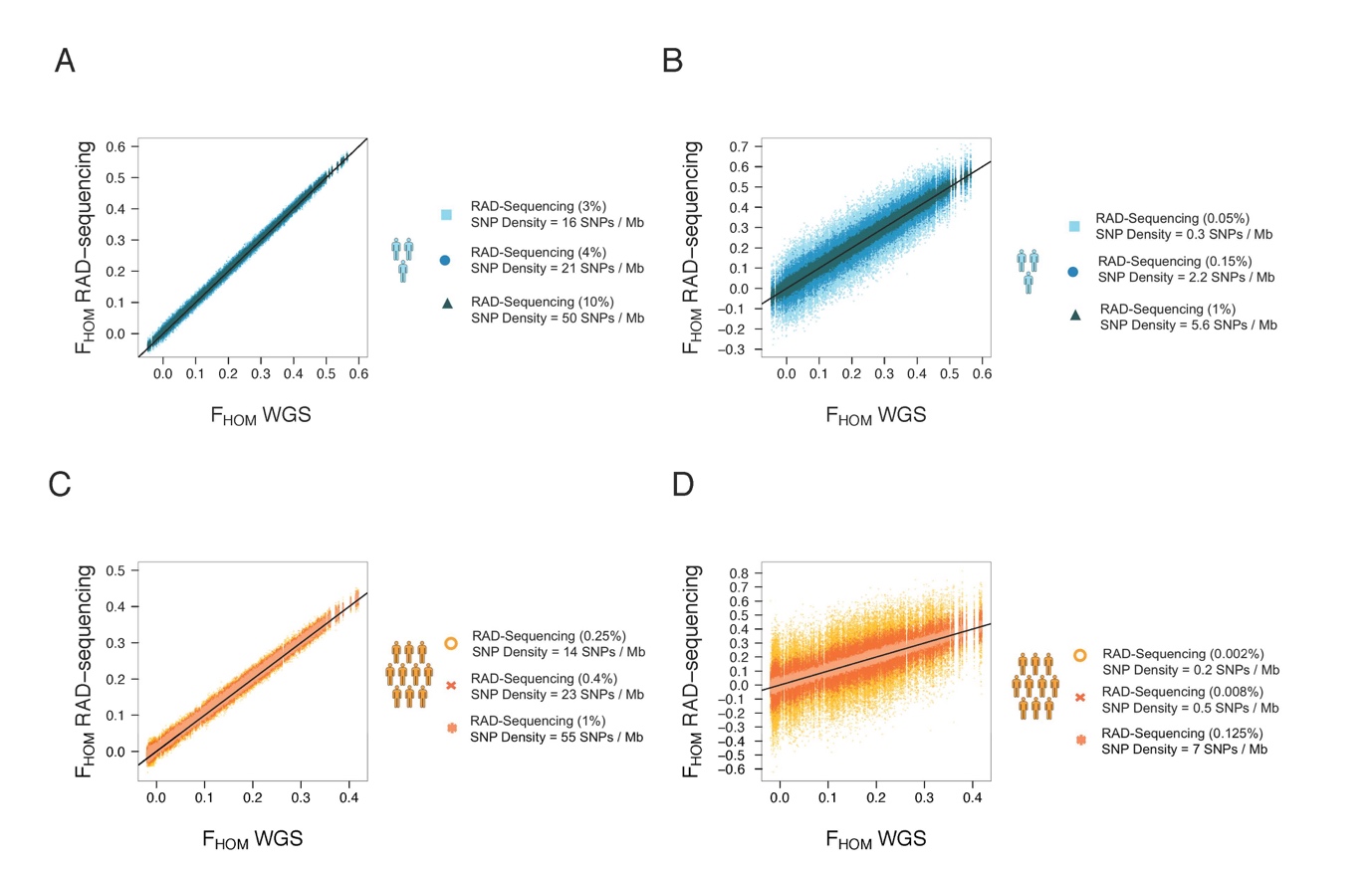

**Figure S10:** Comparison between F_HOM_ estimated with subsampled data mimicking RAD-sequencing (y axis) according to F_HOM_ estimated with WGS data (x axis). **A:** Small population, fraction of genome presented are the same as the fractions of genomes used for ROHs analyses with PLINK (minimum size 100KB) and RAD-sequencing in Figure 1A ; **B:** Small population, fraction of genome presented are the same as the fractions of genomes used for ROHs analyses with RZooRoH and RAD-sequencing in Figure 1B ; **C:** Large population, fraction of genome presented are the same as the fractions of genomes used for ROHs analyses with PLINK (minimum size 100KB) and RAD-sequencing in Figure 1A ; **D:** Small population, fraction of genome presented are the same as the fractions of genomes used for ROHs analyses with PLINK (minimum size 100KB) and RAD-sequencing in Figure 1B.

**Table S1:** Summary of the relationship between inbreeding coefficients (F) estimated with a subsample of the genome and with all the genome (WGS) for: i) F_ROH_ estimated with PLINK: F_ROH PLINK_, ii) F_ROH_ estimated with RZooRoH: F_ROH RZooRoH_ and iii) F_HOM_. For each replicate, the correlation is estimated as the Pearson correlation between F_subsample_ and F_WGS_ and the slope and intercept are extracted from the linear model: $F_{subsample} \sim F_{WGS}$. This table presents the mean (±sd) among replicates.

|  | **% genome sequenced** | **F estimate** | **Correlation ± sd** | **Slope ± sd** | **Intercept ± sd** |
| --- | --- | --- | --- | --- | --- |
| 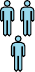 |  |  |  |  |  |
|  | 0.05% | F_ROH RZooRoH_ | 0.929 ± 0.015 | 1.141 ± 0.053 | -0.184 ± 0.021 |
|  |  | F_HOM_ | 0.903 ± 0.022 | 0.995 ± 0.056 | 0.005 ± 0.014 |
|  | 0.15% | F_ROH RZooRoH_ | 0.977 ± 0.005 | 1.143 ± 0.031 | -0.171 ± 0.013 |
|  |  | F_HOM_ | 0.965 ± 0.008 | 0.995 ± 0.031 | 0.006 ± 0.008 |
|  | 1% | F_ROH RZooRoH_ | 0.997 ± 0.001 | 1.117 ± 0.011 | -0.118 ± 0.005 |
|  |  | F_HOM_ | 0.994 ± 0.001 | 0.994 ± 0.013 | 0.006 ± 0.003 |
|  | 3% | F_ROH PLINK_ | 0.337 ± 0.115 | 0.007 ± 0.004 | 0 ± 0 |
|  |  | **F_HOM_** | **0.998 ± 0** | **0.994 ± 0.007** | **0.006 ± 0.002** |
|  | 4% | F_ROH PLINK_ | 0.986 ± 0.004 | 0.733 ± 0.050 | -0.055 ± 0.005 |
|  |  | **F_HOM_** | **0.999 ± 0** | **0.994 ± 0.006** | **0.006 ± 0.002** |
|  | 10% | F_ROH PLINK_ | 0.999 ± 0 | 1.012 ± 0.005 | -0.046 ± 0.002 |
|  |  | **F_HOM_** | **0.999 ± 0** | **0.994 ± 0.004** | **0.006 ± 0.001** |
| 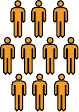 |  |  |  |  |  |
|  | 0.002% | F_ROH RZooRoH_ | 0.702 ± 0.066 | 1.044 ± 0.136 | -0.049 ± 0.042 |
|  |  | F_HOM_ | 0.593 ± 0.084 | 0.988 ± 0.174 | 0.008 ± 0.029 |
|  | 0.008% | **F_ROH RZooRoH_** | **0.916 ± 0.018** | **1.069 ± 0.059** | **-0.063 ± 0.019** |
|  |  | F_HOM_ | 0.857 ± 0.032 | 0.997 ± 0.075 | 0.007 ± 0.013 |
|  | 0.125% | F_ROH RZooRoH_ | 0.996 ± 0.001 | 1.158 ± 0.014 | -0.117 ± 0.004 |
|  |  | F_HOM_ | 0.988 ± 0.003 | 0.992 ± 0.020 | 0.007 ± 0.003 |
|  | 0.25% | F_ROH PLINK_ | 0.503 ± 0.101 | 0.012 ± 0.004 | 0 ± 0 |
|  |  | **F_HOM_** | **0.994 ± 0.001** | **0.993 ± 0.013** | **0.007 ± 0.002** |
|  | 0.4% | F_ROH PLINK_ | 0.995 ± 0.002 | 0.826 ± 0.022 | -0.010 ± 0.001 |
|  |  | **F_HOM_** | **0.996 ± 0.001** | **0.993 ± 0.011** | **0.007 ± 0.002** |
|  | 1% | F_ROH PLINK_ | 0.999 ± 0 | 0.983 ± 0.003 | -0.001 ± 0 |
|  |  | **F_HOM_** | **0.998 ± 0** | **0.992 ± 0.007** | **0.007 ± 0.001** |

**Figure S11**

**Figure S11:** Distributions of the difference in inbreeding rank between WGS and SNP arrays in both populations. ROHs were called with PLINK, minimum size 100KB. Individual inbreeding rank were estimated for the sequencing techniques and the difference between both was obtained for each individual in each subsampling replicate. The distribution of these differences is represented in this figure. **A:** small population and small array; **B:** small population and large array; **C:** large population and small array **D:** large population and large array.

**Figure S12**

**Figure S12:** Distributions of the difference in inbreeding rank between WGS and SNP arrays in both populations. ROHs were called with RZooRoH, 3 HBD classes. Individual inbreeding rank were estimated for the sequencing techniques and the difference between both was obtained for each individual in each subsampling replicate. The distribution of these differences is represented in this figure. **A:** small population and small array; **B:** small population and large array; **C:** large population and small array **D:** large population and large array.

**Figure S13**

**Figure S13**: F_ROH_ estimated with various **percentages of SNPs** (≠ genome) subsampled in the small population and compared to F_ROH_ estimated with WGS data. ROHs were called with PLINK (minimum size = 100KB).

**Figure S14**

**Figure S14**: F_ROH_ estimated with various **percentages of SNPs** (≠ genome) subsampled in the large population and compared to F_ROH_ estimated with WGS data. ROHs were called with PLINK (minimum size = 100KB).

**Figure S15**

**Figure S15:** F_ROH_ estimated with various **percentages of SNPs** (≠ genome) subsampled in the small population and compared to F_ROH_ estimated with WGS data. ROHs were called with RZooRoH (3 HBD classes).

**Figure S16**

**Figure S16:** F_ROH_ estimated with various **percentages of SNPs** (≠ genome) subsampled in the large population and compared to F_ROH_ estimated with WGS data. ROHs were called with RZooRoH (3 HBD classes). No ROHs calling with RZooRoH with a random subsampling with more than 20% of the SNPs were performed in the large population due to computational constrains.

**Figure S17**

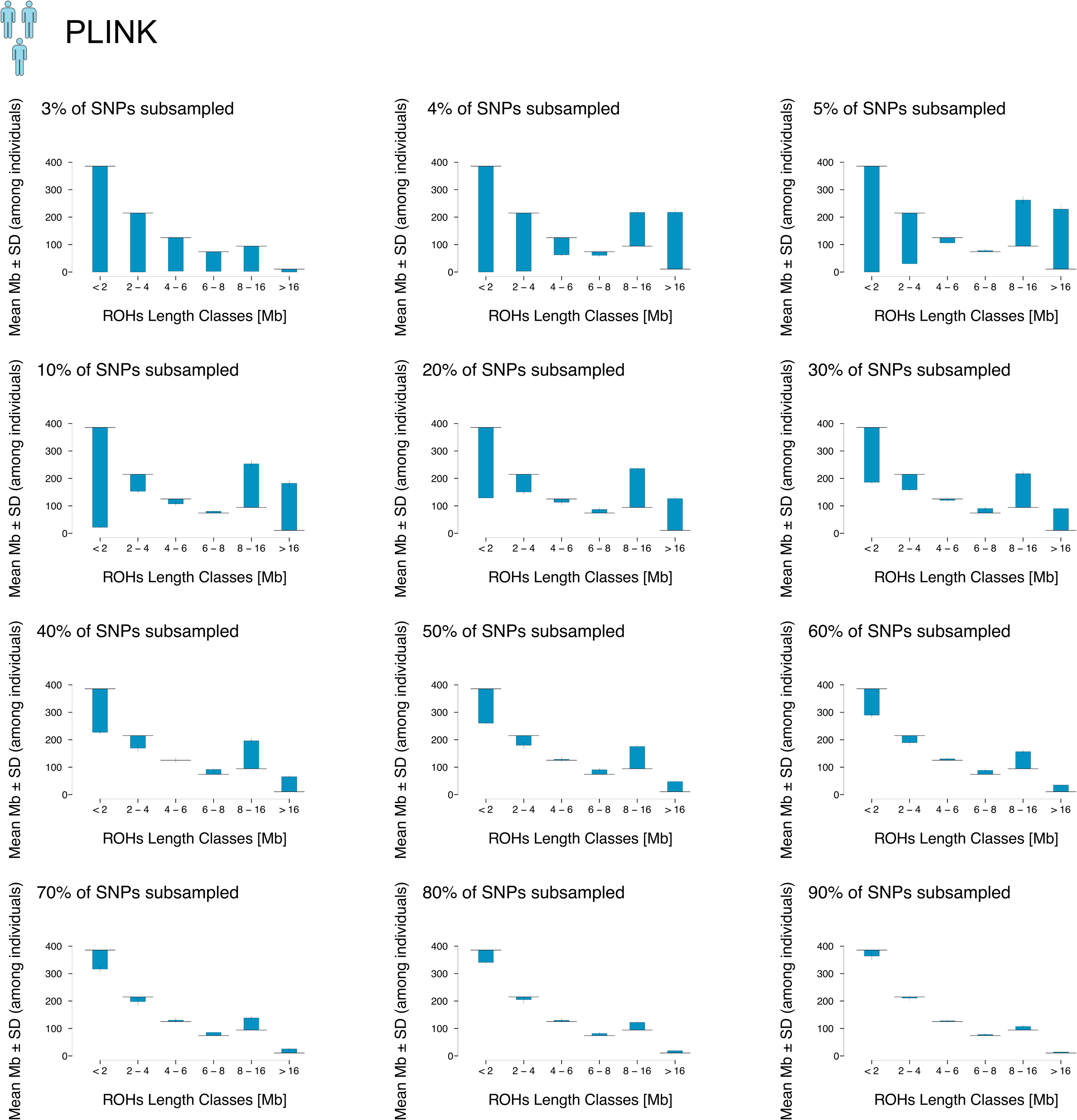

**Figure S17:** Comparison of ROHs distributions between WGS (black continuous line) and various percentages of SNPs subsampled (grey dashed line) in the small population. ROHs were called with PLINK, minimum size 100KB.

**Figure S18**

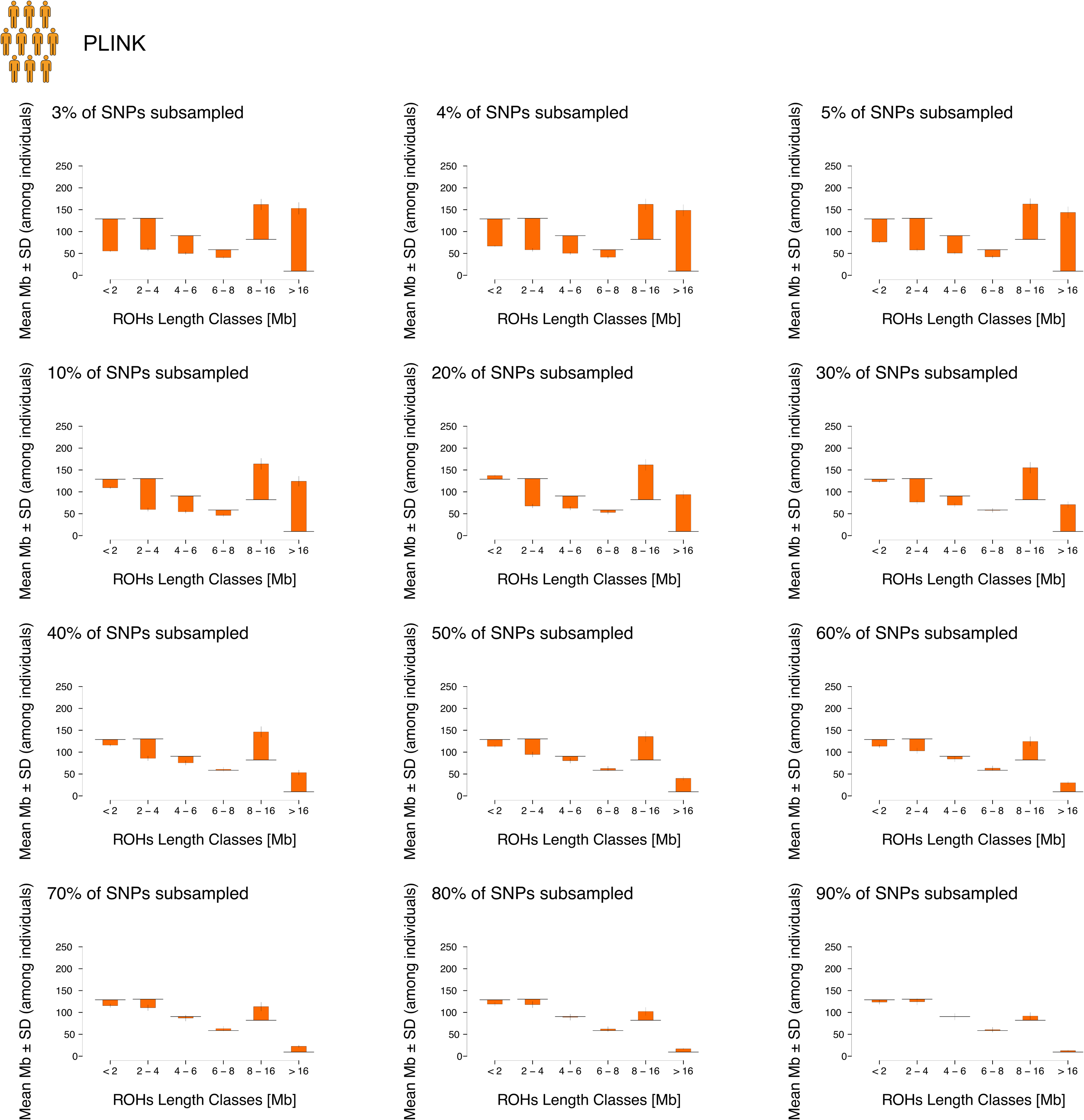

**Figure S18:** Comparison of ROHs distributions between WGS (black continuous line) and various percentages of SNPs subsampled (grey dashed line) in the large population. ROHs were called with PLINK, minimum size 100KB.

**Figure S19**

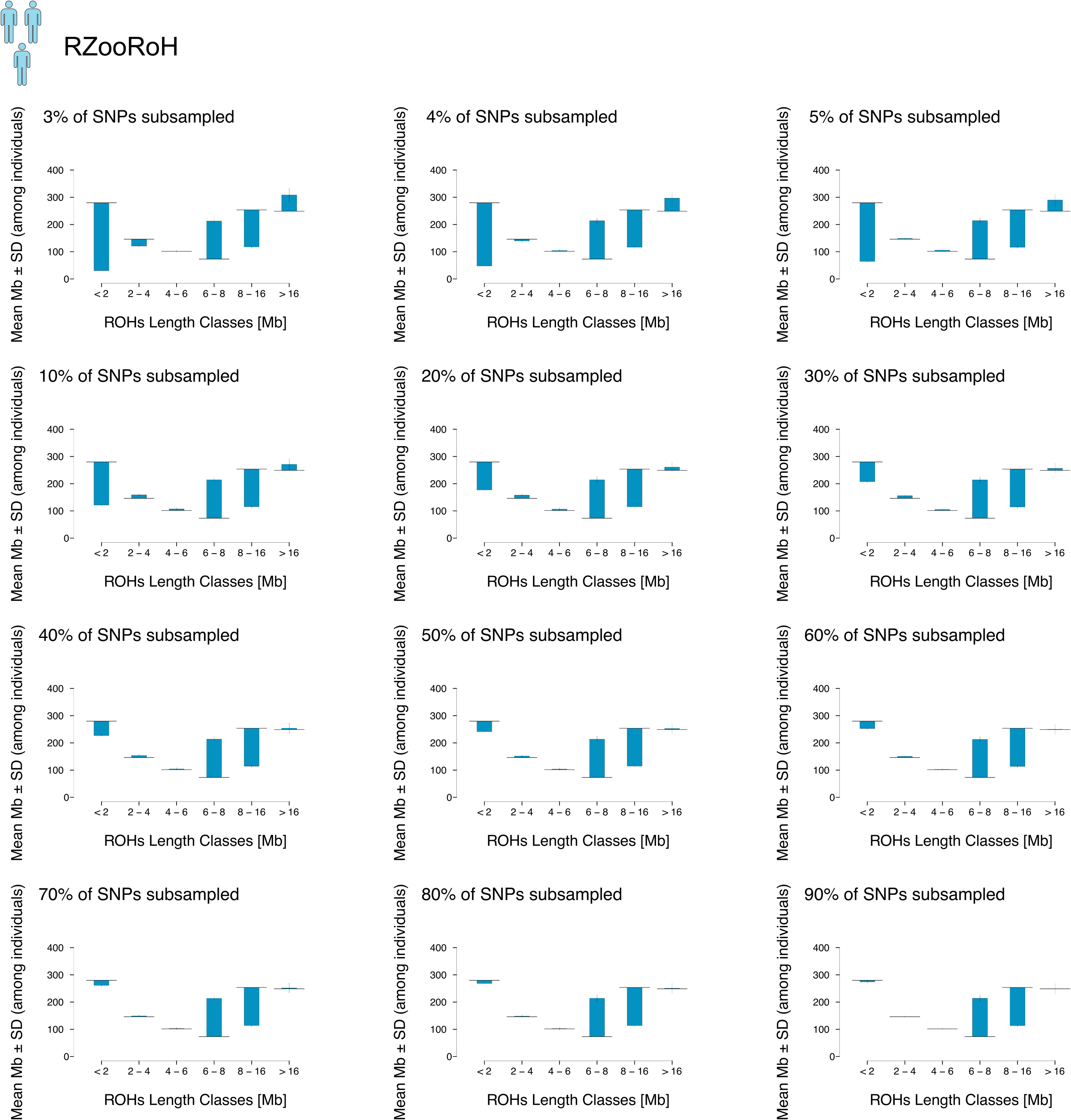

**Figure S19**: Comparison of ROHs distributions between WGS (black continuous line) and various percentages of SNPs subsampled (grey dashed line) in the small population. ROHs were called with RZooRoH (3 HBD classes).

**Figure S20**

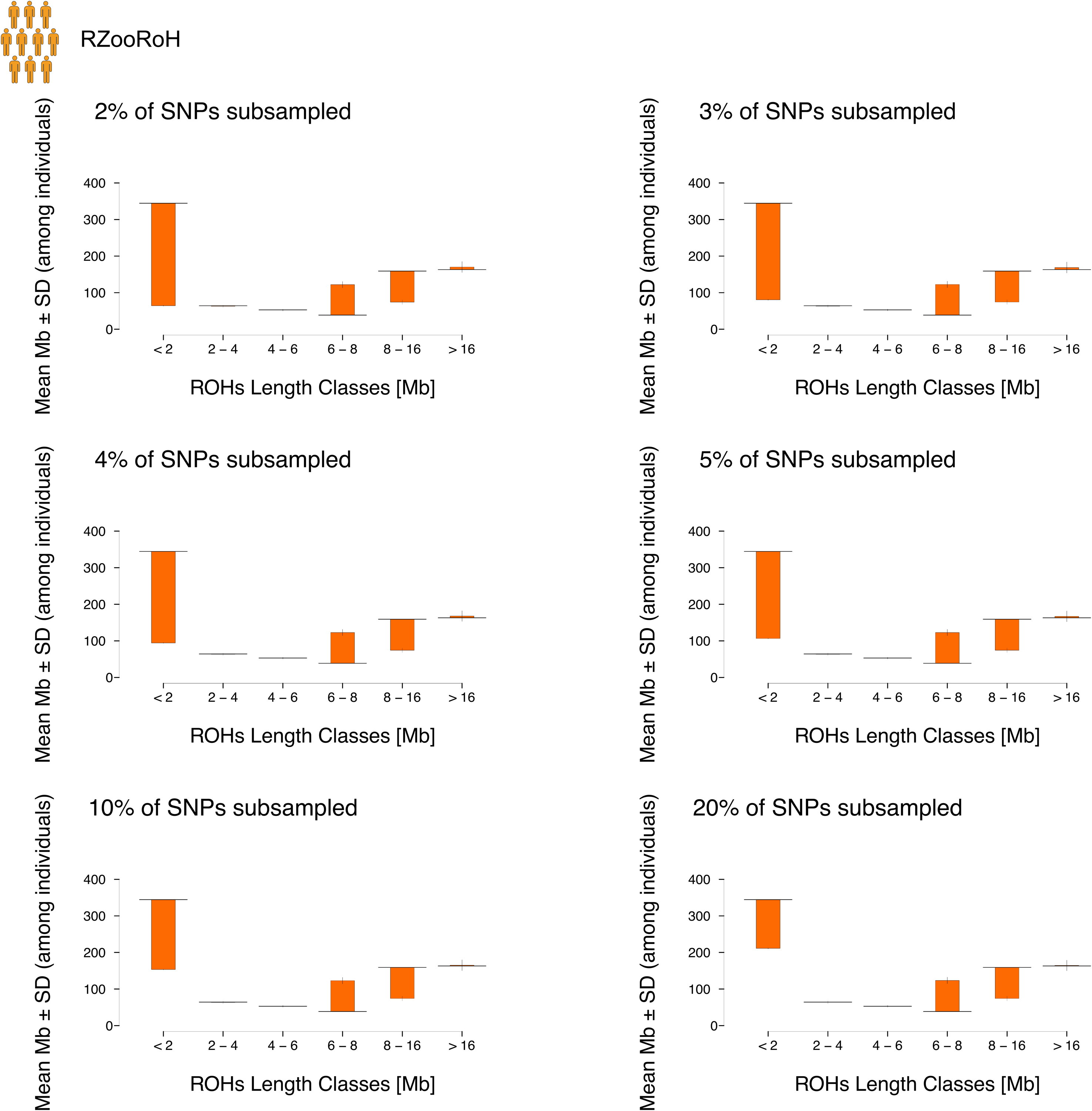

**Figure S20:** Comparison of ROHs distributions between WGS (black continuous line) and various percentages of SNPs subsampled (grey dashed line) in the small population. ROHs were called with RZooRoH (3 HBD classes). No ROHs calling with RZooRoH with a random subsampling with more than 20% of the SNPs were performed in the large population due to computational constrains.

**Figure S21**

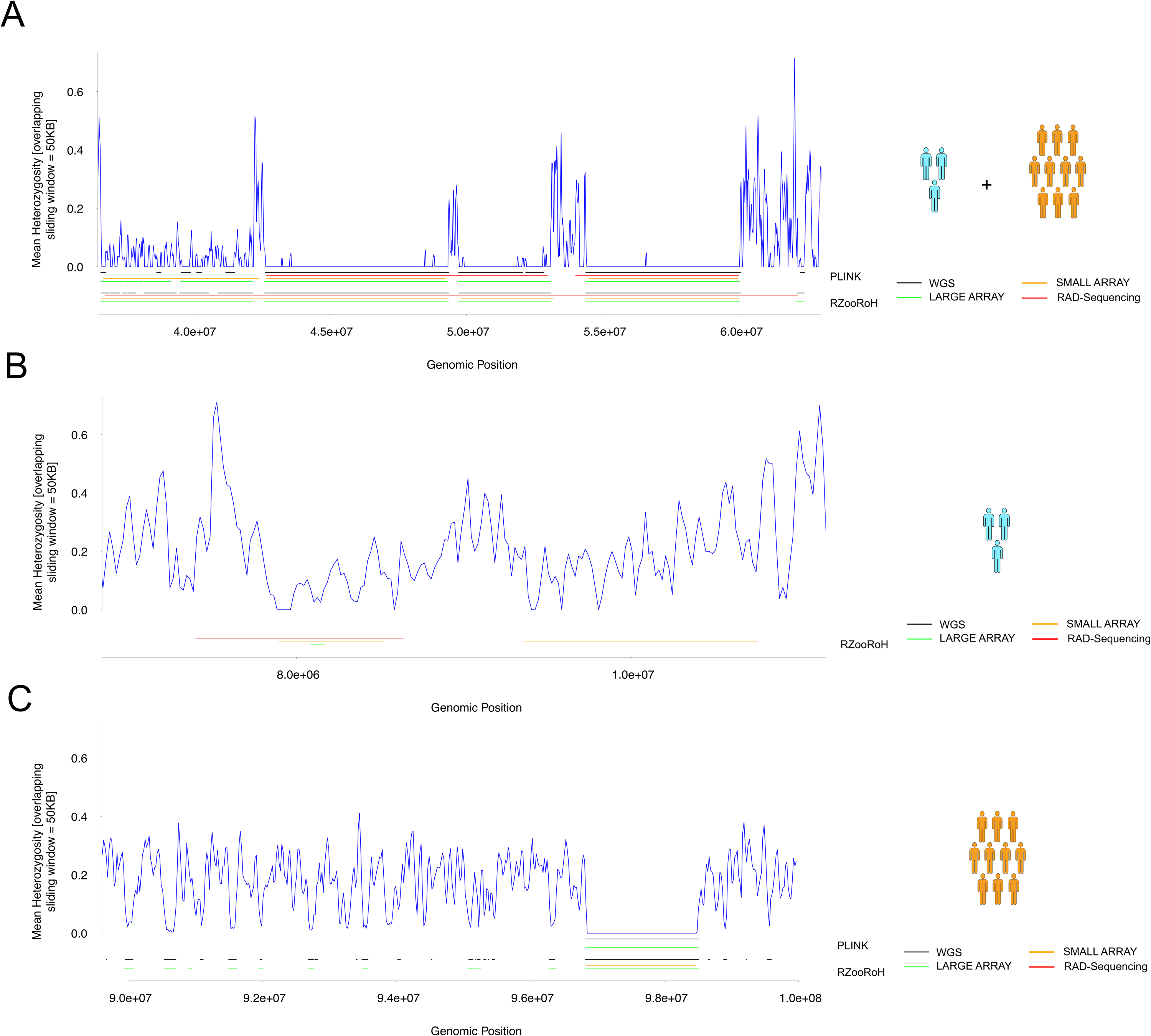

**Figure S21:** Mean Heterozygosity (y axis) according to genomic position (x axis). One example of ROH detection in one subsampling replicate of one simulation replicate is plotted below for both ROHs calling methods. **A:** Example of reduced genomic representation merging small adjacent ROHs into a larger one. **B:** Example of spurious ROH detection with reduced genomic representation and RZooRoH. **C:** Example of many small ROHs in the genome in the large population. Only the large array allowed to detect these small ROHs.

**Figure S22**

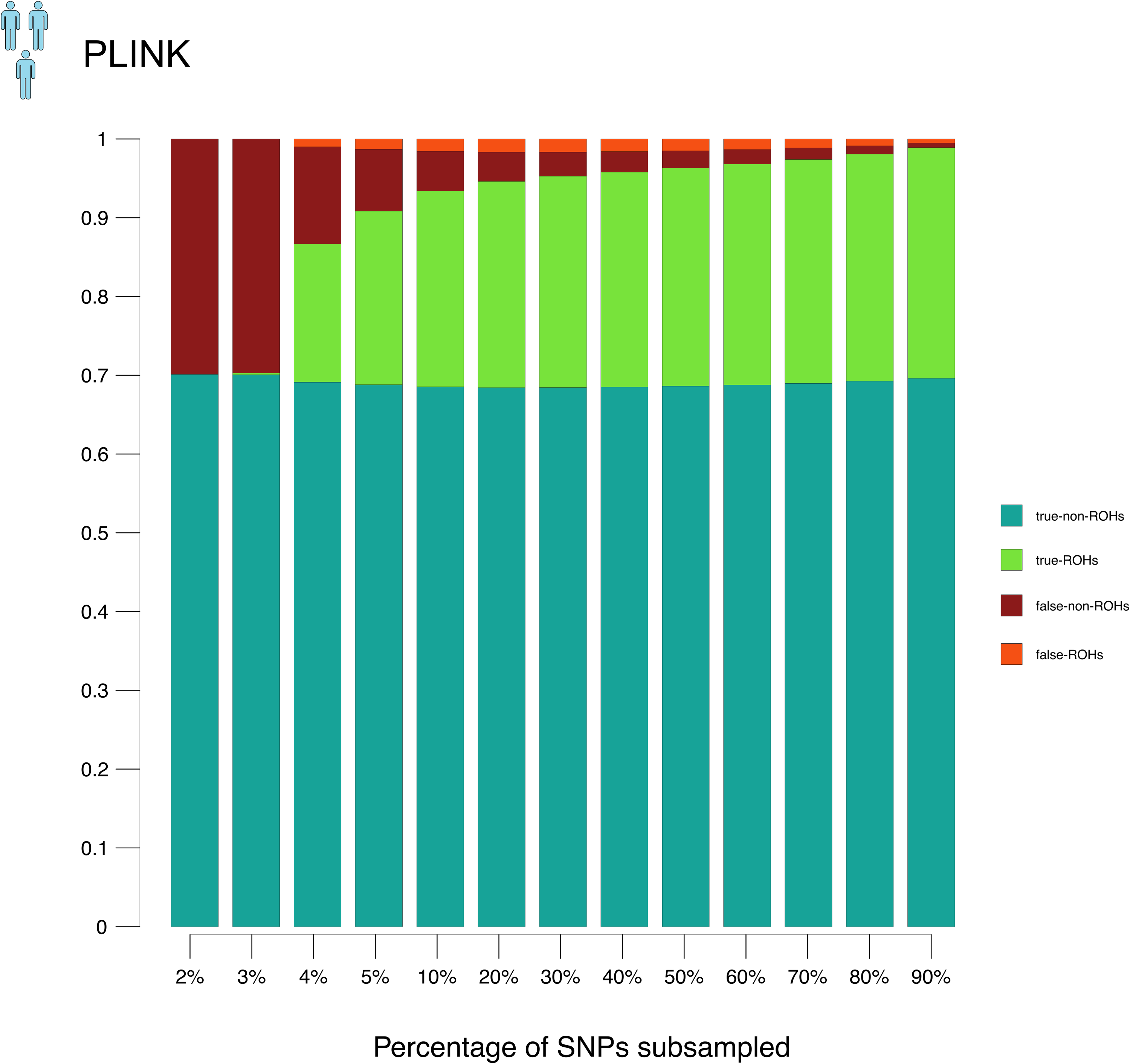

**Figure S22:** Fraction of genome correctly and incorrectly assigned within and outside ROHs for the small population. ROHs were called with PLINK, minimum size = 100KB.

**Figure S23**

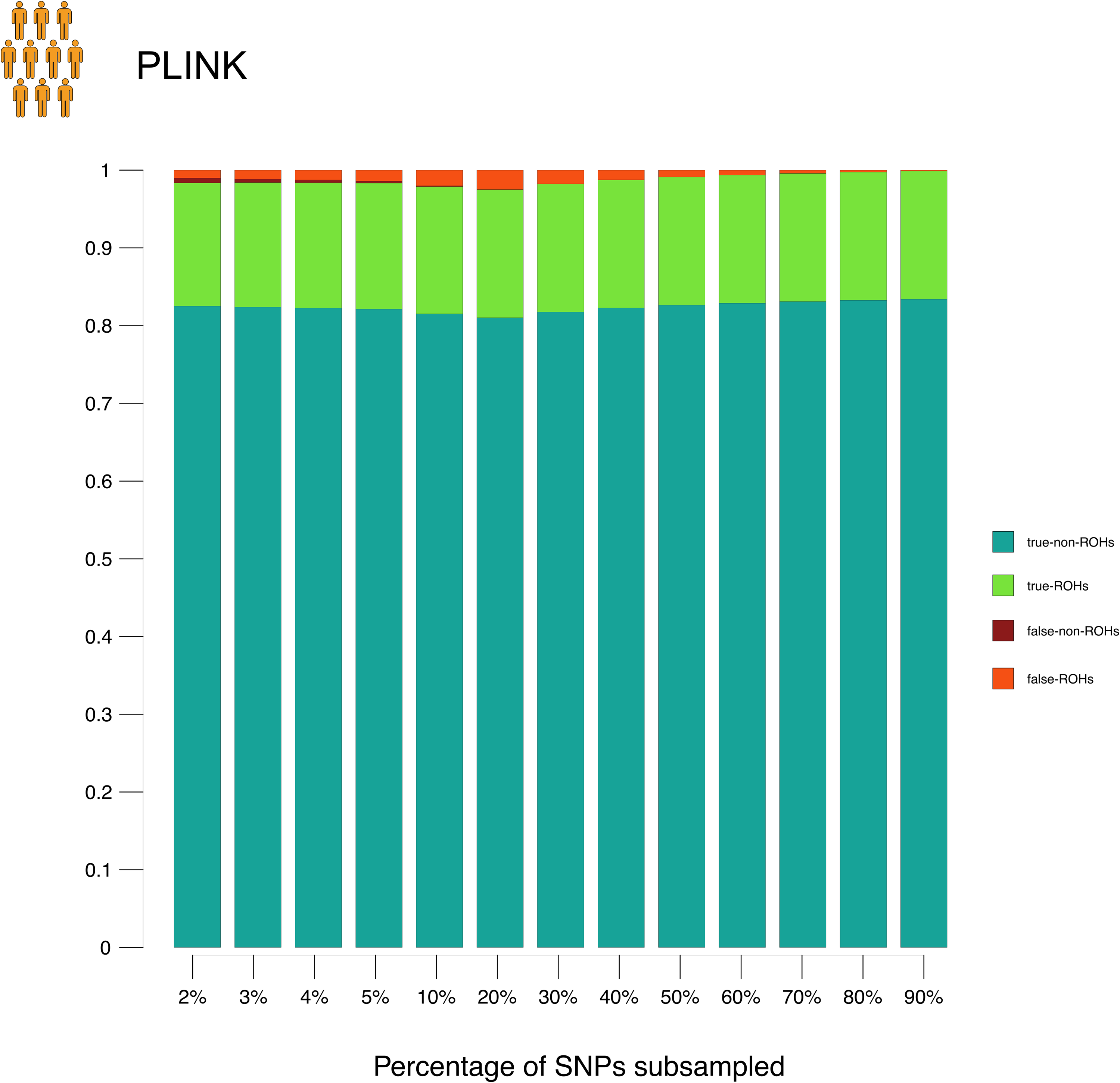

**Figure S23:** Fraction of genome correctly and incorrectly assigned within and outside ROHs for the small population. ROHs were called with PLINK, minimum size = 100KB.

**Figure S24**

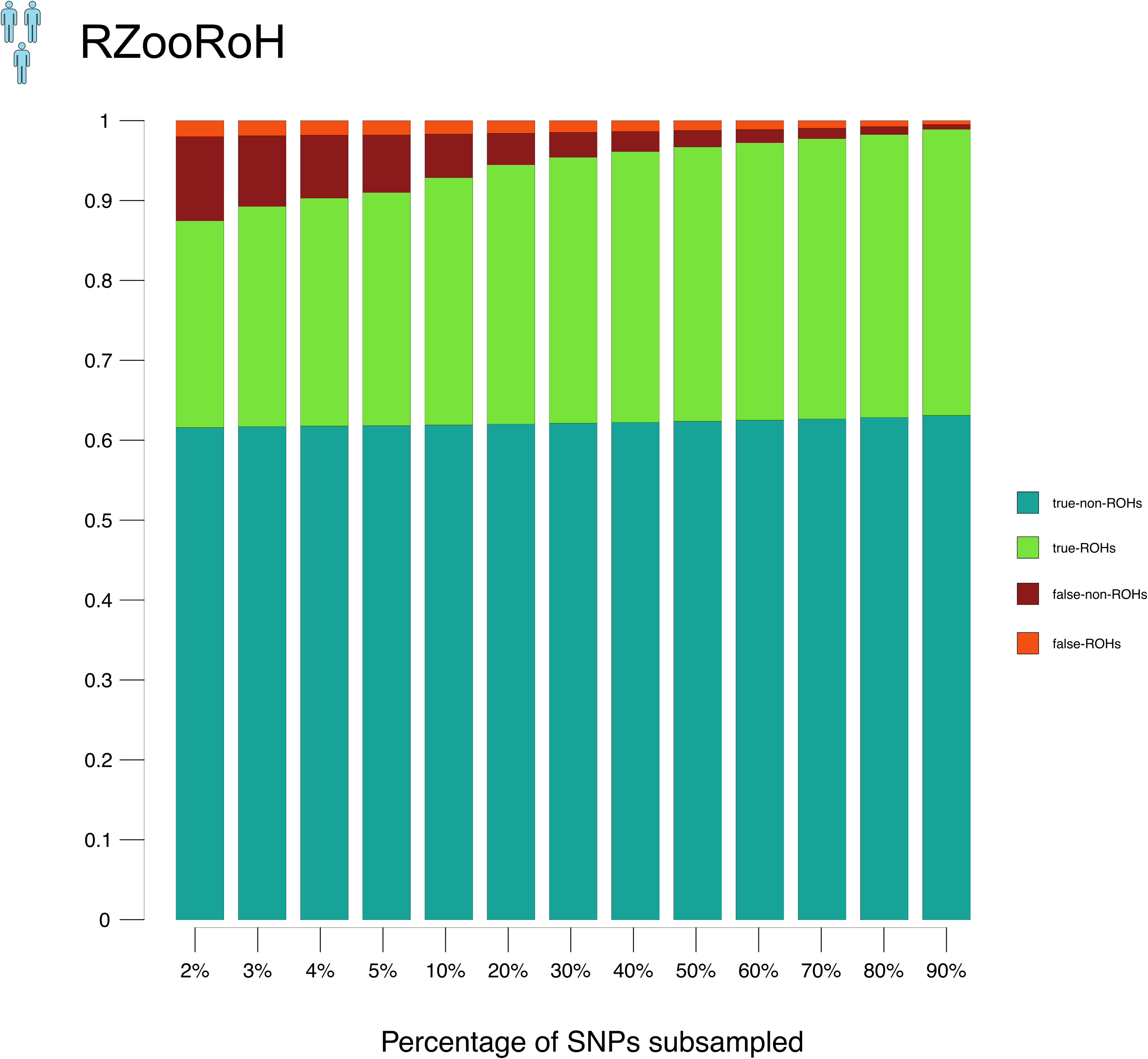

**Figure S24:** Fraction of genome correctly and incorrectly assigned within and outside ROHs for the small population. ROHs were called with RZooRoH (3 HBD classes).

**Figure S25**

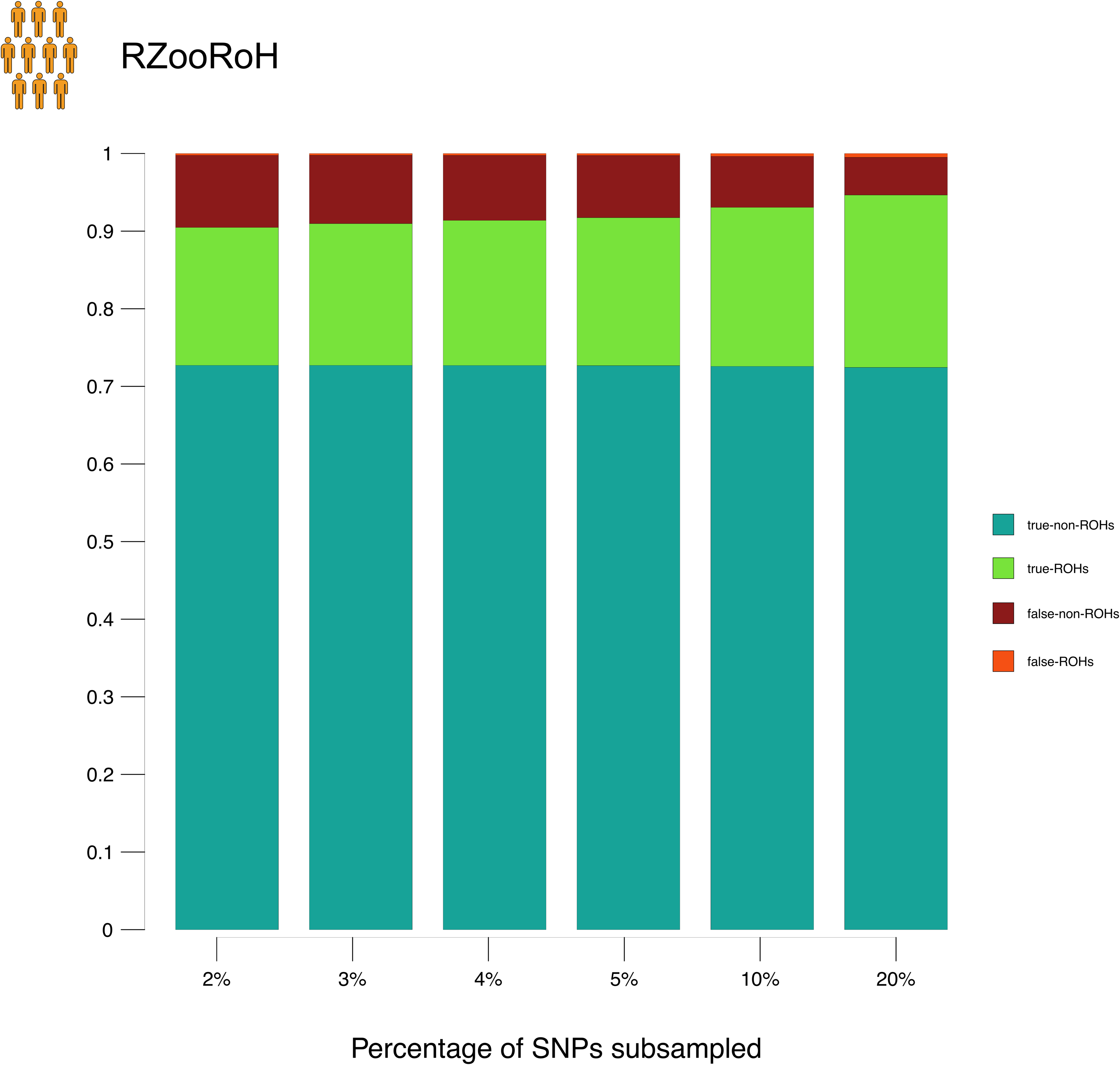

**Figure S25:** Fraction of genome correctly and incorrectly assigned within and outside ROHs for the small population. ROHs were called with RZooRoH (3 HBD classes). No ROHs calling with RZooRoH with a random subsampling with more than 20% of the SNPs were performed in the large population due to computational constrains.

**Figure S26**

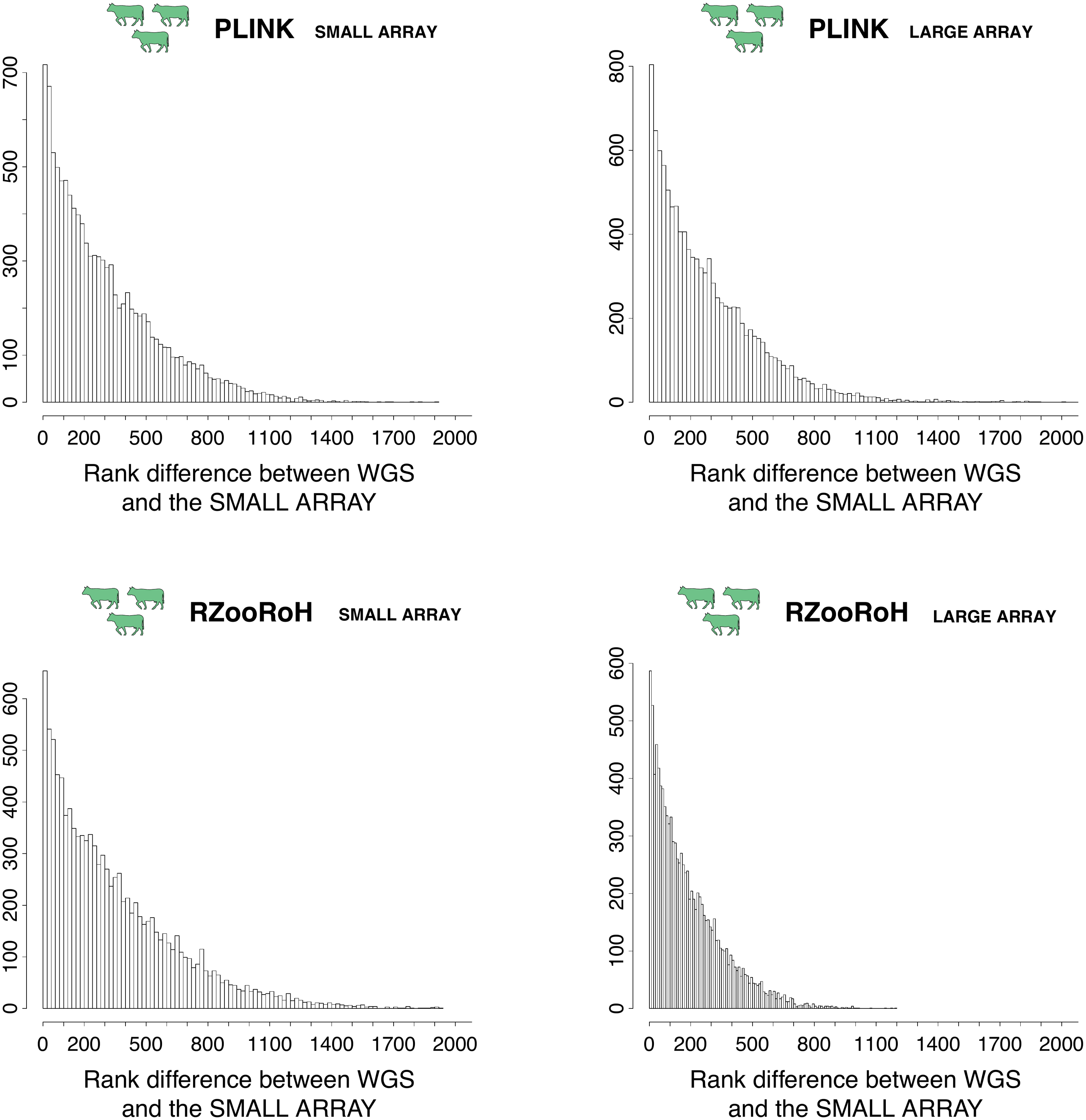

**Figure S26:** Distributions of the difference in inbreeding rank between WGS and SNP Arrays in the cattle population. ROHs were called with both PLINK, minimum size 100KB and RZooRoH, 3 HBD class model. Individual inbreeding rank were estimated for both sequencing techniques and the difference between both was obtained for each individual. The distribution of these differences is represented in this figure.
